# A Conserved Geometric Code: Extracellular Matrix Curvature Directs Cell Migration Strategy via Nuclear Mechanosensing

**DOI:** 10.64898/2026.03.24.713851

**Authors:** Bo Cheng, Yan Liu, Zhao Xu, Huan Gao, Qihui Sun, Lingwenyao Kong, Xi Wang, Ningman Dai, Longji Dong, Fanyu Li, Kunnuo Yu, Caiyun Wang, Lin Wang, Yuan Li, Min Lin, Ting Wen, Tian Jian Lu, Ye Li, Feng Xu

## Abstract

Cells navigate complex tissue microenvironments defined by intricate physical cues, yet how they interpret the three-dimensional geometry of the extracellular matrix (ECM) remains an open question. Current models often fail to account for the tortuous architectures found in physiological tissues. Here, we demonstrate that ECM curvature functions as a tissue-specific geometric code read by the cell nucleus. By mapping collagen architectures across cancers and tissues, we find unique curvature fingerprints preserved during metastasis. Using micro-engineered substrates, we show that high curvature imposes localized nuclear bending stress, triggering a Lamin A/C-cPLA2-Ca^2+^ mechanotransductive cascade. This sensor rewires the cytoskeleton from longitudinal stress fibers to a cortical actomyosin network, driving a sharp transition from fast mesenchymal migration to a slower, exploratory amoeboid phenotype. We term this “nuclear curvotaxis”, establishing a physical principle linking static geometry to dynamic strategy, with implications for predicting metastatic risk, understanding immune exclusion, and designing bio-instructive scaffolds for tissue engineering.

## INTRODUCTION

Living cells are not passive passengers within tissues; they are active readers of a complex physical language written in the extracellular matrix (ECM)^1–3^. This physical microenvironment regulates essential behaviors, including differentiation, proliferation, and migration, through dynamic mechanotransduction pathways^4–8^. Traditionally, physical cues are well established as guidance factors across biology, *i.e.*, matrix stiffness (gradient sensing known as durotaxis) and fiber alignment (contact guidance along ECM fibres). It is well established that cells migrate directionally up gradients of stiffness (durotaxis) and align their trajectories along linearized collagen fibers (contact guidance or topotaxis)^4,9^. In the context of cancer, the stiffening of the tumor stroma and the radial alignment of collagen fibers, known as Tumor-Associated Collagen Signatures (TACS), are recognized as potent drivers of metastasis and predictors of poor patient survival^10–13^.

Despite these advances, a critical dimension of the ECM remains largely deciphered: its geometry. Native tissues are rarely flat or perfectly linear; they are geometrically complex, non-Euclidean environments characterized by tortuous fibers, undulating interfaces, and intricate pore networks with varying degrees of curvature^14–17^. From the sinusoidal crimp of healthy tendon collagen to the disordered, looped networks of desmoplastic stroma, curvature is a ubiquitous feature of tissue architecture^18–20^. While recent theoretical and in vitro studies have described “curvotaxis”, the preference of cells to migrate based on substrate curvature, the relevance of this phenomenon in the complex 3D physiological environment is poorly understood^7,8^. Specifically, it is unknown whether cells can sense the inherent curvature of an ECM fiber as a distinct cue from its stiffness or density, and how this geometric information is integrated into migratory decision-making.

Navigating this 3D geometry imposes significant physical constraints on the cell, particularly on the nucleus. As the largest and stiffest organelle, the nucleus acts as the rate-limiting factor in confined migration^21–23^. Emerging research has fundamentally shifted our view of the nucleus from a passive cargo to an active mechanosensory. Pioneering studies have established the “nuclear ruler” hypothesis, demonstrating that nuclear deformation in confined spaces triggers a mechanotransduction pathway involving the unfolding of the nuclear envelope (NE), the recruitment of cytosolic phospholipase A2 (cPLA2), and the release of calcium to modulate actomyosin contractility^21,24^. This mechanism allows cells to measure pore size and adapt their passage. However, whether this nuclear sensing mechanism enables cells to detect curvature, to distinguish between a straight narrow channel and a curved one, remains a major knowledge gap.

We hypothesized that ECM curvature functions as a conserved, tissue-specific “geometric code” that instructs cell migration strategy via nuclear mechanosensing. Unlike stiffness, which can be dynamically remodeled by cells, we posit that fiber curvature is a stable structural feature determined by tissue architecture. We further reasoned that the nucleus serves as the primary “curvature sensor”, transducing geometric bending into a binary switch between migration modes. This study integrates quantitative pathology, micro-engineering, and single-cell transcriptomics to test this hypothesis.

In this paper, we identify ECM curvature as a robust predictor of cell behavior. We show that diverse human and murine tissues possess unique “curvature fingerprints” that are preserved even at metastatic sites. Using engineered microenvironments that recapitulate these fingerprints, we demonstrate that high curvature acts as a physical checkpoint, forcing cells to switch from a fast, persistent, mesenchymal-like migration mode to a slow, exploratory, amoeboid-like phenotype. Mechanistically, we reveal that this switch is driven by “nuclear curvotaxis”: high curvature imposes localized bending stress on the nuclear envelope, activating a Lamin A/C-cPLA2-Ca²⁺ signaling axis that rewires the cytoskeleton from stress fibers to a cortical actomyosin network. This mechanism is distinct from canonical YAP/TAZ mechanotransduction, highlighting the specificity of the nuclear tension pathway. Our findings not only provide a new framework for understanding metastatic dissemination and immune cell infiltration but also offer novel geometric parameters for the design of bio-instructive materials and mechanomedicine therapeutics.

## RESULTS

### ECM curvature is a conserved geometric fingerprint of each tissue, remaining stable across disease progression

To establish the geometric context in which cells operate *in vivo*, we first mapped the collagen fiber curvature in a variety of pathological tissues. We analyzed histological sections from human breast tumors, oral fibrotic lesions (periodontitis), skin scars, and melanomas, using a custom image analysis pipeline to quantify fiber curvature at each point in the ECM (**Supplementary Fig. S1** and **Fig. S2**). Masson’s trichrome staining of breast cancer specimens revealed dense collagen bundles encasing tumour nests (**Fig. 1A**, top), with intensity mapping showing maximal fibre density at tumour margins and gradual decline towards peripheral tissue (**Fig. 1A**, middle). Orientation analysis indicated highly aligned fibres near the tumour edge, transitioning to heterogeneous orientations with distinct curved segments at boundary zones (**Fig. 1A**, bottom). Fibre orientation heatmaps further highlighted this gradient: fibres near the tumour displayed uniform orientation angles, whereas distal regions exhibited heterogeneous angle distributions with distinct curved segments emerging at boundary zones (**Fig. 1B**). These curved fibres disrupted linear arrays and implied localized mechanical heterogeneity, potentially influencing cell trajectory patterns.

**Figure 1.**
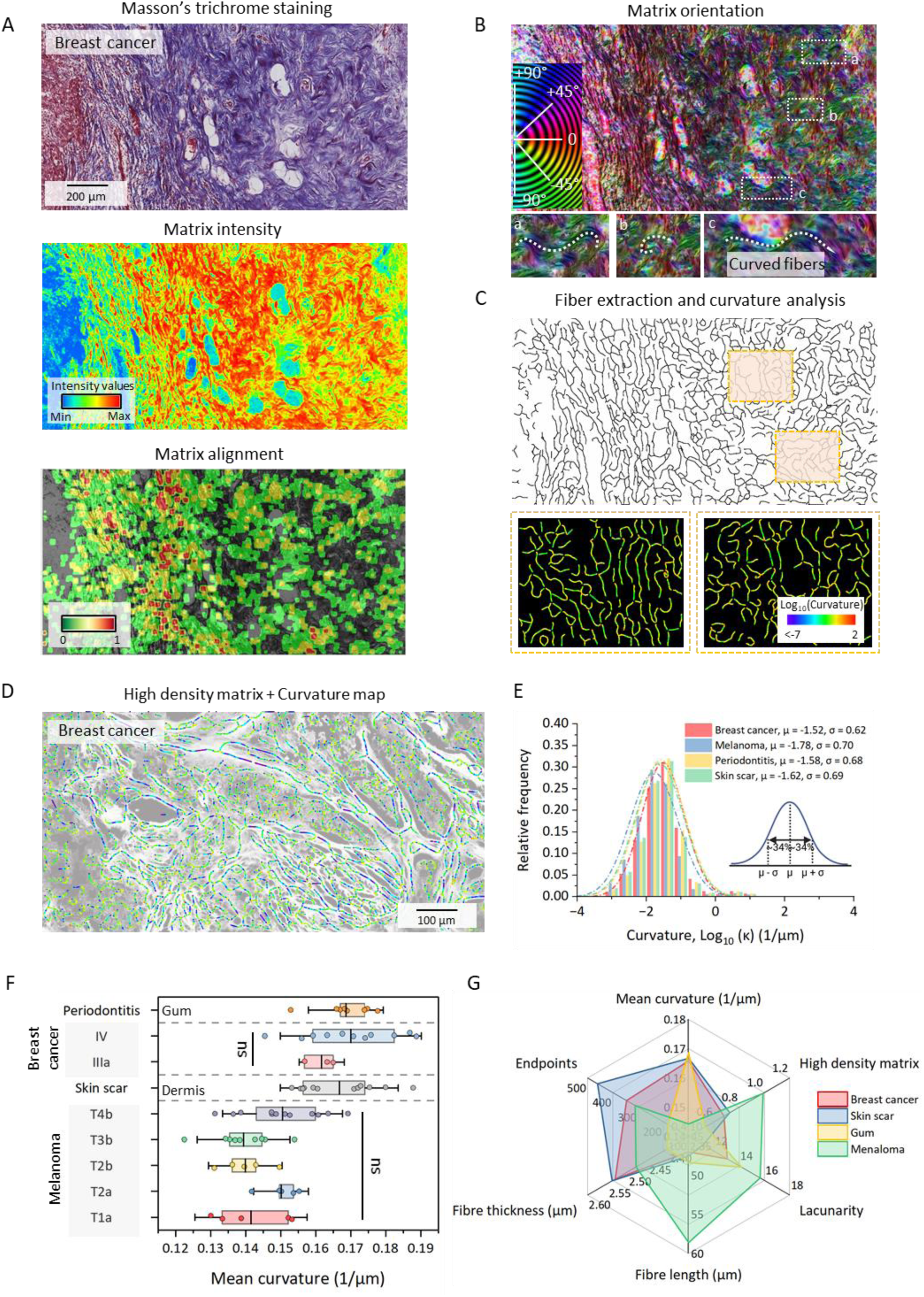
Mean ECM curvature is a conserved geometric fingerprint across tissues. **A,** Breast cancer ECM architecture. Top: Masson’s trichrome shows dense collagen bundles (blue/red) surrounding tumor nests. Middle: ECM intensity heatmap (optical density) with red/yellow for high density, blue for low. Bottom: ECM alignment map: warm colors indicate parallel fibres (alignment index ∼1), gray indicates low parallelism. **B,** Orientation analysis using OrientationJ colour-codes fibre angles from −90° (blue) to +90° (red). Near the tumour, uniform colours indicate parallel arrays, whereas peripheral zones display heterogeneous orientations with curved segments that locally disrupt linear organization (insets). **C,** Automated ECM extraction skeletonises the collagen network (top) and computes point-wise curvature (bottom). Curvature heatmaps (log₁₀ scale) identify regions enriched in high curvature (>2 μm⁻¹; warm colours) and low curvature (<1 μm⁻¹; cool colours). **D,** Overlay of curvature and density maps in breast cancer ECM shows that high-density, aligned fibre regions coincide with low curvature (blue), whereas boundary zones, where density decreases and orientation shifts, are enriched in high-curvature segments (red/orange). **E,** Curvature histograms and Gaussian fits for breast cancer, melanoma, periodontitis and skin scar ECM. Each tissue exhibits a distinct mean curvature (*μ*) and spread (*σ*); μ is lowest in breast cancer and highest in melanoma, with intermediate values in periodontitis and scar. **F,** Mean curvature is conserved within a given tissue type across pathology stages and anatomical sub-sites. While *μ* differs between tissues, it is statistically unchanged across progression stages within the same tissue, indicating that curvature is set early in remodelling and remains stable. Melanoma T1a, n = 5; Melanoma T2a, n = 5; Melanoma T2b, n = 5; Melanoma T3b, n = 9; Melanoma T4b, n = 12; Skin scar, n = 12; Breast cancer IIIa, n = 3; Breast cancer V, n = 11; Periodontitis, n = 9. **G,** Radar plots of six ECM metrics, mean curvature, high-density matrix fraction, fibre endpoints, mean fibre thickness, mean fibre length and lacunarity, reveal multi-parameter structural “fingerprints” for each tissue. Breast cancer ECM is dense, aligned and low curvature with few endpoints; melanoma and periodontitis are more porous and curved; scar ECM features thicker, longer fibres. Scale bars, 200 μm (a, top), 50 μm (b, insets), 100 μm (d). Statistical tests and non-significant (NS) differences are indicated in panels.

To quantify these geometric features, we skeletonized the ECM network and calculated point-wise curvature (**Fig. 1C**). Curvature maps revealed that low-curvature fibres predominated in high-density, highly aligned regions near tumours, whereas high-curvature fibres were enriched in areas where density dropped and orientation shifted (**Fig. 1D** overlay). This spatial association suggests curvature changes coincide with ECM structural transitions, possibly marking sites of altered tensile stress or matrix remodelling.

We extended this quantitative analysis to multiple pathological tissues, breast cancer, melanoma, periodontitis, and skin scar, constructing curvature histograms for each (**Supplementary Fig. S3** and **Fig. S4**). Across all these diverse tissues, we found that the distribution of fiber curvatures in each tissue follow a roughly Gaussian profile characterized by a tissue-specific mean curvature (**Fig. 1E**). Notably, the average fiber curvature (*μ*) differs between tissue types, for example, collagen in breast tumor stroma is on average less curved (lower *μ*) than in melanoma or scar tissue, yet within a given tissue type the mean curvature is highly consistent across different samples and disease stages (**Fig. 1F**). In breast cancer, for instance, late-stage (invasive) tumors exhibit virtually the same mean ECM curvature as early-stage tumors. Similarly, melanomas from primary to advanced stages maintains a constant characteristic curvature value. This finding indicates that each tissue establishes a particular ECM curvature “set point” early on, which persists despite pathological remodeling. In practical terms, while tissues undergoing disease may see drastic changes in collagen density or alignment, their inherent fiber curvature pattern remains unchanged.

Besides, to capture broader ECM structural context, we profiled six additional fibre metrics: high-density matrix percentage, fibre endpoints, thickness, length, lacunarity, and alignment (**Fig. 1G** and **Supplementary Fig. S5**). Radar plots illustrated distinct multi-parameter “fingerprints”: breast cancer ECM was dense and aligned with low curvature and few endpoints; melanoma and periodontitis appeared more porous and curved; skin scar was characterized by thicker, longer fibres. Together, these findings establish mean ECM curvature as a stable, tissue-specific parameter fixed early in pathological remodelling, implying that local curvature gradients rather than global shifts represent the functionally relevant variation.

### Curvature magnitude acts as a physical switch between fast-persistent and exploratory migration modes

We next asked whether this conserved curvature fingerprint in primary tumors is recapitulated at distant metastatic sites, where migrating cells colonize new microenvironments. To test this, we examined ECM geometry in metastases from two orthotopic cancer models. In a mouse melanoma model (B16F10), dermal tumors were allowed to metastasize to the lungs. We observed that collagen in the lung metastases forms dense arc-shaped fibres surrounding the tumor foci, very much like the curved collagen bundles at the perimeter of the primary melanoma (**Fig. 2A**). Quantitative curvature mapping in these metastatic lesions yields a mean fiber curvature of approximately 0.20 μm⁻¹, closely matching the high-curvature signature of the primary melanoma’s ECM. Similarly, in an orthotopic human breast cancer model (MDA-MB-231 cells), lung metastases exhibit collagen fibres with more gentle bends and a mean curvature around 0.16 μm⁻¹, mirroring the low-curvature ECM fingerprint of the primary breast tumor. (**Fig. 2B**) These data show that migrating cancer cells often encounter similar curvature cues in the metastasis site as in their tissue of origin. It appears that disseminated tumor cells may “carry” their preferred ECM geometry with them, or preferentially thrive in distant niches that incidentally have a similar curvature profile to the primary tumor. In summary, ECM curvature signatures are preserved at metastatic sites, suggesting that stable geometric features of the matrix can be maintained or re-established during metastasis.

**Figure 2.**
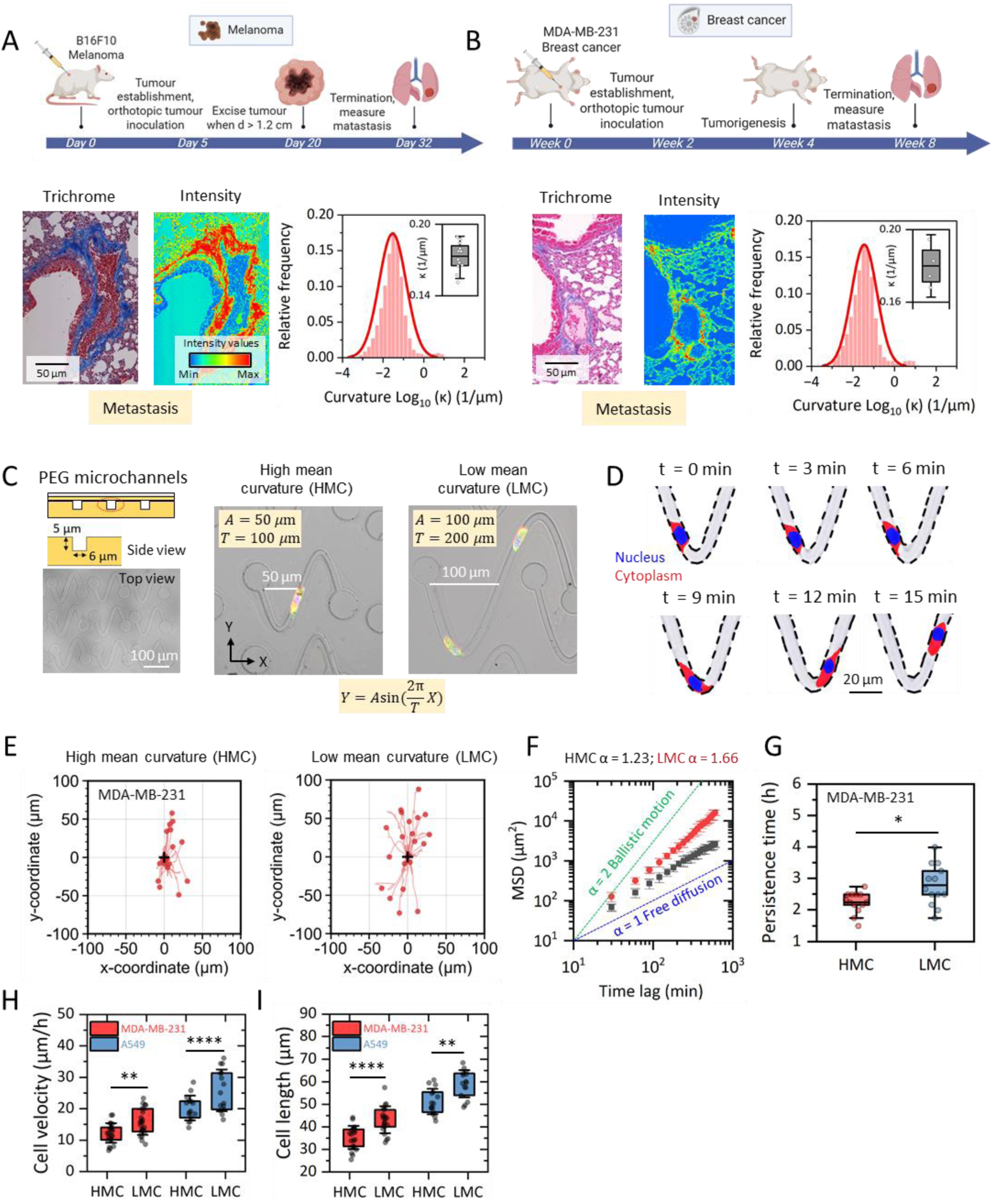
Tissue-specific ECM curvature is preserved at metastatic sites and switches migration modes in engineered confined geometries. **A,** In an orthotopic melanoma model (B16F10), tumours are implanted dermally, resected at >1.2 cm diameter and allowed to form lung metastases. Masson’s trichrome staining of perivascular ECM at metastatic foci reveals dense collagen arcs surrounding tumour clusters. Intensity maps identify high-fibre-density zones at the tumour-host interface, and curvature histograms show a mean curvature μ ≈ 0.20 μm⁻¹, matching the high-curvature fingerprint of primary melanoma stroma. n = 22 samples. **B,** In an orthotopic breast cancer model (MDA-MB-231), primary mammary tumours generate lung metastases by week 8. Collagen bundles at metastatic margins exhibit more gently curved trajectories. Curvature mapping yields μ ≈ 0.16 μm⁻¹, corresponding to the low-curvature ECM characteristic of breast tumour stroma. n = 4 samples. **C,** Sinusoidal PEG microchannels are designed to reproduce these tissue curvature fingerprints: high mean curvature (HMC) channels (*A* = 50 μm, *T* = 100 μm) and low mean curvature (LMC) channels (*A* = 100 μm, *T* = 200 μm). Schematics show top view, cross-section and sinusoidal wall equations. **D,** Time-lapse imaging of cells migrating in curved channels (nucleus, blue; cytoplasm, red) reveals distinct behaviours. Positions at 3 min intervals illustrate cell trajectories along successive bends. **E,** Representative migration paths of MDA-MB-231 cells in HMC (left) and LMC (right) microchannels. Trajectories in LMC channels are longer and straighter, with greater net displacement. **F,** Mean squared displacement (MSD) analyses show scaling exponents (α) approaching ballistic motion (α ≈ 1.66) in LMC channels and reduced exponents (α ≈ 1.23) in HMC channels, consistent with a switch from persistent to exploratory motion. **G,** Directional persistence times are significantly higher in LMC than in HMC channels for MDA-MB-231 cells, indicating enhanced guidance along low-curvature tracks. HMC, n = 13; LMC, n = 13. **H,** Instantaneous migration velocities for three cancer cell lines are consistently higher in LMC than in HMC channels, demonstrating that low curvature supports faster movement across lineages. MDA-MB-231, n = 23; A549 = 13. **I,** Cells in LMC channels are more elongated along the migration axis than in HMC channels, aligning morphology with persistent trajectories. MDA-MB-231, n = 23; A549 = 13. Scale bars, 50 μm (A,B, histology), 100 μm (C, top view), 20 μm (D). Box plots show mean, quartiles and range; significance levels and test types are indicated; ns, not significant.

Based on our tissue measurements, we designed controlled *in vitro* microenvironments to directly test how curvature influences cell migration behavior. We fabricated sinusoidal microchannels from biocompatible hydrogels, tuned to have either high mean curvature (HMC, with amplitude *A* = 50 μm, period *T* = 100 μm, peak *κ*∼0.20 μm⁻¹) or low mean curvature (LMC, with amplitude *A* = 100 μm, period *T* = 200 μm, peak *κ*∼0.16 μm⁻¹) along their wavy walls (**Supplementary Fig. S6A-B** and **Fig. 2C**), values corresponding to the *in vivo* ECM curvatures. Apart from geometry, all other physical parameters of the channels (such as stiffness and confinement width) were kept identical, so that any differences in cell behavior could be attributed to curvature alone (**Supplementary Fig. S6C-D**). We then seeded various cell types (including MDA-MB-231 breast carcinoma cells and A549 lung carcinoma cells) into these channels and tracked their migration using time-lapse microscopy (**Fig. 2D**).

Strikingly, we observed a sharp phenotypic switch in how cells migrated depending on the curvature of their confinement. In LMC channels, migration trajectories were long, straight and rapid (**Fig. 2E**); mean squared displacement (MSD) slopes approached ballistic motion (α ≈ 1.66; **Fig. 2F**), persistence times were significantly higher than in HMC channels (**Fig. 2G**), and instantaneous velocities exceeded those in HMC in MDA-MB-231 breast carcinoma cells (**Fig. 2H**). Similarly to the 2*A* = *T* channel, cells in the high-curvature region (with an amplitude of *A* = *T*) also exhibited reverse-direction or loop-back migration, which is rarely observed in the low-curvature region (**Supplementary Fig. S7A-B**). Morphologically, LMC cells were elongated along the migration axis and have more protrusions (**Fig. 2I** and **Supplementary Fig. S7C-D**). These curvature-dependent migration phenotypes were reproduced in A549 lung carcinoma cells (**Supplementary Fig. S8**), demonstrating that low curvature geometries act as efficient “guidance tracks” enabling fast, highly persistent migration, whereas high curvature promotes exploratory modes with diminished net displacement. By matching in-vitro channel curvature to in-vivo ECM fingerprints, these assays directly link native tissue geometry to functional migration behaviour.

### Curvature amplifies mechanosensing pathways and expands invasion-prone cell subpopulations

To examine whether curvature also reprograms transcriptional states relevant to migration and invasion, we performed single-cell RNA sequencing (scRNA-seq) on MDA-MB-231 cells grown under three ECM geometries: High curvature modelled with polyacrylamide (PA) microsphere substrates of radius 40 µm (***R40***); Low curvature with PA microsphere of radius 115 µm (***R115***); and flat control, *i.e.*, planar PEG hydrogels (**Fig. 3A-B**).

**Figure 3.**
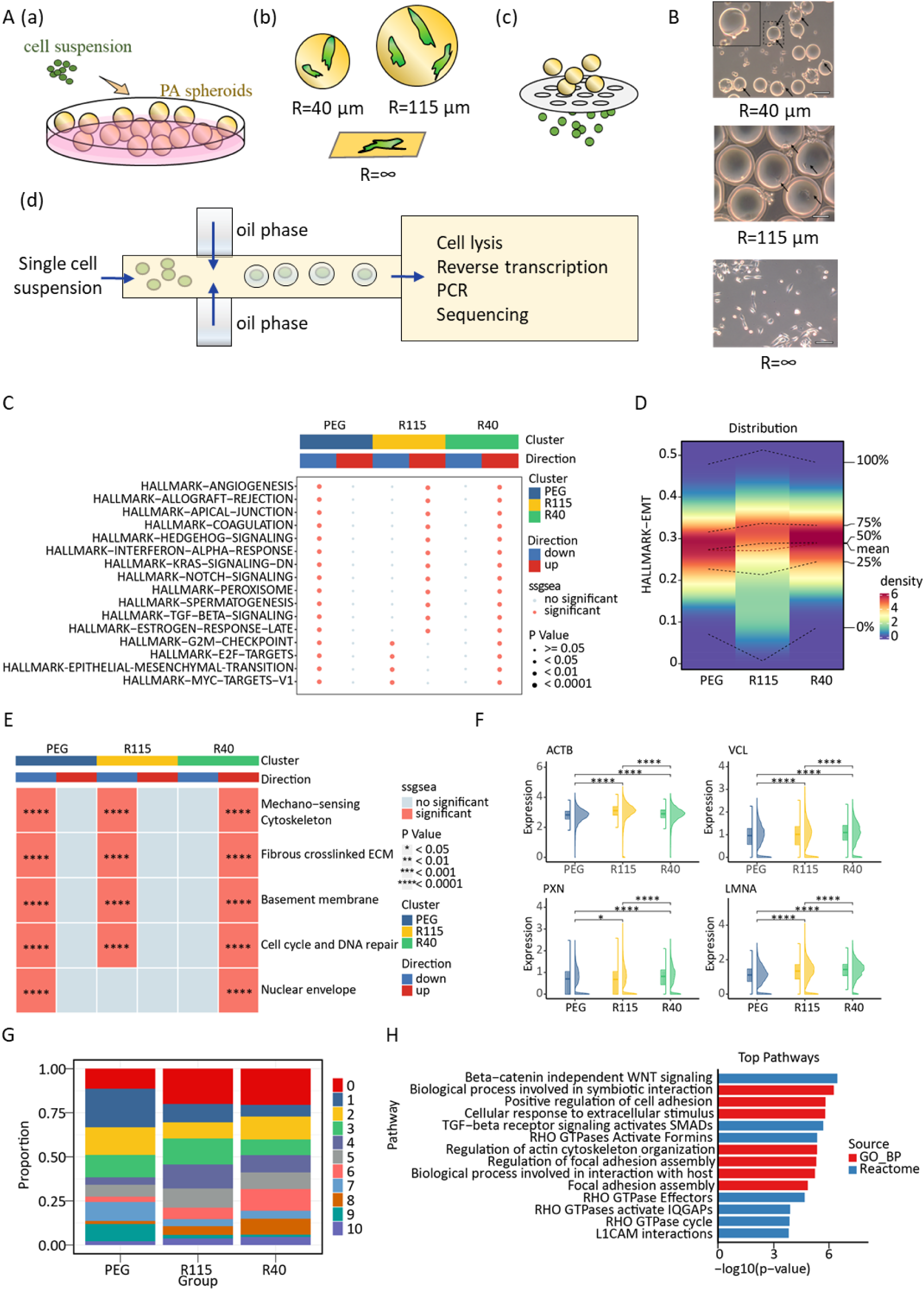
ECM curvature activates mechanosensitive transcriptional programs and expands invasion-prone epithelial subpopulations. **A,** Workflow for curvature-controlled single-cell RNA sequencing (scRNA-seq). Cell suspensions form spheroids in PEG microwells, which are transferred onto collagen-coated polyacrylamide substrates with defined curvature radii (R40, high curvature; R115, low curvature; flat control). Spheroids are then dissociated into single cells, captured in a microfluidic device, and processed for scRNA-seq. **B,** Brightfield images of spheroids on substrates of increasing curvature illustrate distinct morphologies and attachment patterns. **C,** Gene set variation analysis (irGSEA package, ssGSEA score) of hallmark pathways shows progressive enrichment of EMT and migration-associated programs with increasing curvature, peaking under R40 (high curvature) conditions. **D,** Distribution of HALLMARK–EMT scores reveals a curvature-dependent shift: EMT activity is lowest on flat substrates, intermediate on R115 and highest on R40. **E,** Curvature drives strong activation of mechanosensing and structural programs. ssGSEA scores for cytoskeleton organization, fibrous crosslinked ECM, basement membrane, DNA repair and nuclear envelope pathways are maximal under high curvature. **F,** Violin plots show upregulation of core mechanosensitive genes (VCL, PXN, LMNA, ACTB) in high-curvature conditions, consistent with reinforced focal adhesion, lamina and actin networks. **G,** Stacked bar plots of epithelial subcluster composition demonstrate curvature-dependent remodelling of cell states, with specific clusters expanding under high curvature. **H,** Pathway analysis of these curvature-expanded subclusters reveals enrichment of ECM regulation, RHO GTPase signalling and cytoskeletal remodelling, and identifies populations with reduced differentiation potential and enhanced invasive capacity.

Transcriptomic analysis revealed that increasing curvature leads to progressive enrichment of specific gene programs related to cell migration and mechanosensing. Using differential expression analysis (DEA),weighted gene co-expression network analysis (WGCNA), and functional enrichment analysis, we observed a graded upregulation of hallmark pathways as we moved from flat, to LMC, to HMC conditions (**Supplementary Figs. S9A-C**). In particular, cells in the HMC environment show strong activation of epithelial-mesenchymal transition (EMT) gene signatures, cytoskeletal remodeling pathways, ECM adhesion and crosslinking pathways, and nuclear envelope/lamina regulation programs, compared to flat culture (**Fig. 3C-D**, **Supplementary Fig. S9D-E**). These transcriptional changes are consistent with heightened mechanosensing demands and structural adaptation to a curved environment (**Supplementary Fig. S9F-G**). Notably, focal adhesion genes like vinculin (VCL) and paxillin (PXN), nuclear lamina gene LMNA, and actin gene ACTB are all significantly upregulated in high curvature conditions (**Fig. 3E-F, Supplementary Fig. S9H**), aligning with the idea that cells reinforce their adhesion and nuclear structure to cope with curvature-induced stresses. In contrast, cells on flat or low-curvature substrates maintain comparatively lower expression of these mechano-responsive genes.

Clustering the single-cell transcriptomes uncovers distinct cell subpopulations whose abundance depends on curvature (**Fig. 3G; Supplementary Figs. S10A-C**). Under high curvature, we found an expansion of certain clusters enriched for Rho GTPase signaling, actomyosin contractility, and matrix-degradative pathways (**Fig. 3H**). These clusters also exhibit lower differentiation scores by CytoTRACE analysis, suggesting a more progenitor-like, stem-like state. In other words, curvature selectively expands a subpopulation of cells with traits of high motility and invasiveness, reminiscent of mesenchymal or leader-cell phenotypes (**Supplementary Figs. S10D-E**). In contrast, low-curvature conditions do not strongly favor these invasive clusters; instead, a more differentiated or epithelial-like population dominated. This indicates that curvature not only acutely influences migration behavior, but can also reshape the phenotypic composition of a cell population over time, potentially priming cells for invasion or metastasis.

Integrating these results, we arrive at a cohesive model: high-curvature environments activate broad mechanotransductive and migratory gene programs (EMT-like changes) in cells and enrich for aggressive, less-differentiated cell subpopulations; yet paradoxically, those same conditions lead to physically slower and less directed migration because the cells are constantly reorienting and experiencing nuclear stress. Low-curvature environments impose less transcriptional “strain” on cells, they do not strongly induce EMT or stress pathways, but they provide an optimal geometric architecture for efficient migration, allowing even partially mesenchymal cells to migrate fast and straight. In essence, ECM curvature can modulate both the immediate mechanics of cell migration and the long-term phenotypic state of cells. We term this concept “cellular curvature memory”, wherein exposure to certain geometric curvature landscapes can leave cells in a more invasive state even after they exit that environment. These findings underscore that geometric cues like curvature are integrated into cellular regulatory networks, linking the physical microenvironment to gene expression and cell state in a manner relevant to disease progression (for example, a tumor region with high ECM curvature might biochemically prime cells for invasion even if their net movement is slowed).

### High curvature confinement induces pronounced nuclear envelope bending and a distinct high-curvature nuclear morphology

Because the nucleus is a large object that must deform to accommodate surrounding geometry, we reasoned that nuclear morphology could provide a direct readout of curvature sensing. We developed an imaging approach to segment each nucleus into sub-regions corresponding to the local curvature of the microchannel wall. In cells migrating in sinusoidal channels, we defined the “bend region” (BR) of the nucleus as the portion in contact with the maximally curved part of the channel wall (the inner curve of a wave), and the “side region” (SR) as the portion along a relatively straighter segment of the channel (the outer flank of a wave) (**Fig. 4A**). This intra-nuclear mapping allowed us to compare nuclear shape metrics between high-curvature versus low-curvature zones within the same cell and across different curvature conditions.

**Figure 4.**
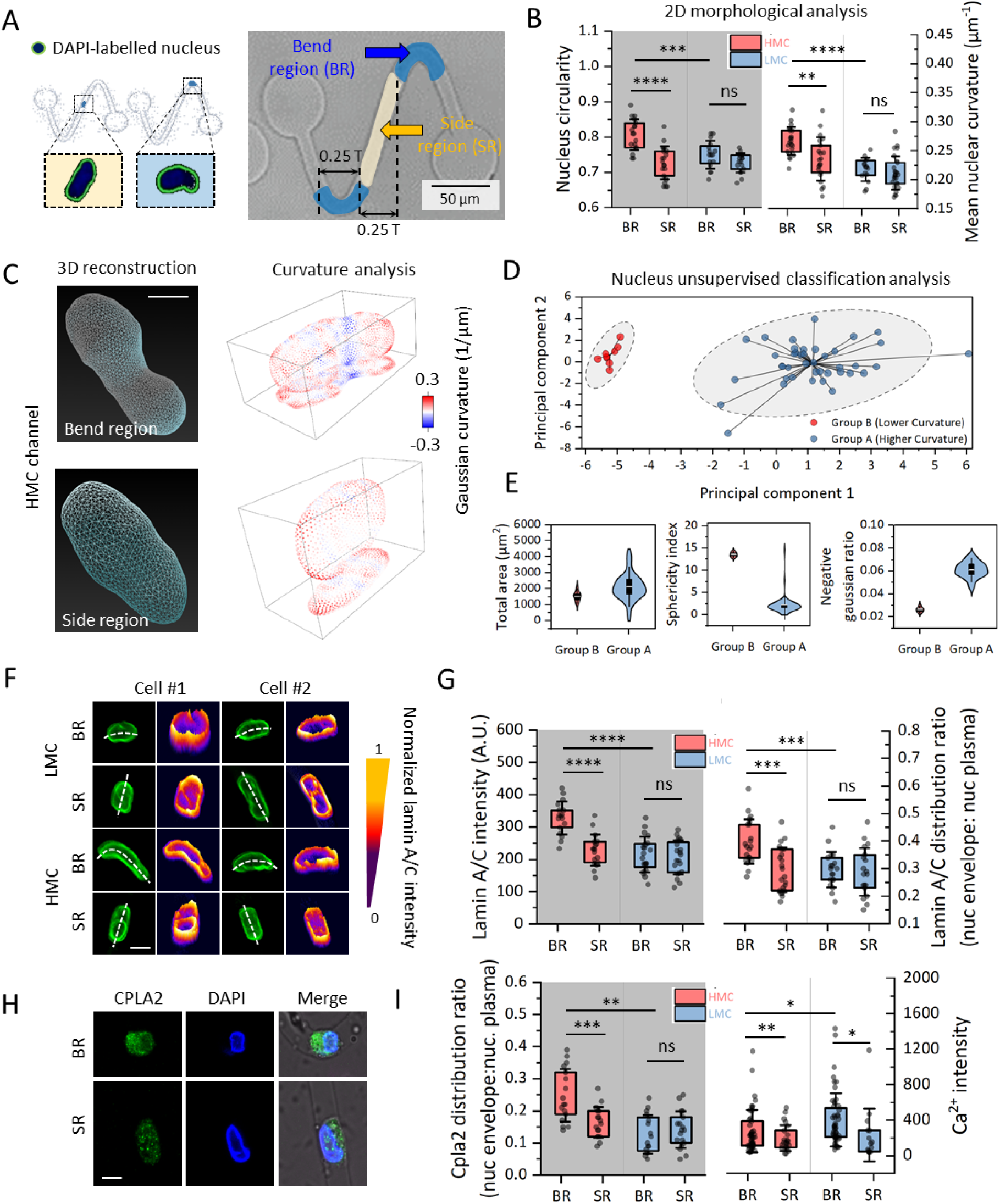
High ECM curvature induces local nuclear bending, morphometric reclassification and nuclear mechanotransduction. **A,** Nuclei of cells migrating in sinusoidal channels are segmented into bend regions (BR, adjacent to local wall curvature peaks) and side regions (SR, along straighter channel sections). **B,** Quantification shows significantly reduced circularity and increased mean nuclear curvature in BR nuclei under high mean curvature (HMC) compared to SR nuclei and to nuclei in low mean curvature (LMC) channels. Circularity: HMC-BR, n = 18; HMC-SR, n = 21; LMC-BR, n = 16; LMC-SR, n = 17. Curvature: HMC-BR, n = 22; HMC-SR, n = 21; LMC-BR, n = 15; LMC-SR, n = 26. **C,** Three-dimensional reconstructions and Gaussian curvature maps reveal alternating positive and negative curvature patches along BR nuclear envelopes in HMC channels, indicating complex folding driven by curved confinement, whereas SR and LMC nuclei remain comparatively smooth. **D,** Principal component analysis of 18 curvature–morphometry variables followed by k-means clustering (*k* = 2) delineates two nuclear classes: Group A (high bending) and Group B (low bending). BR nuclei from HMC channels predominantly populate Group A. **E,** Group A nuclei display lower sphericity and altered area relative to Group B, confirming that HMC confinement generates a distinct high-curvature morphometric state. **F,** Immunostaining for Lamin A/C shows local enrichment at BR regions of bent nuclei. Dashed lines denote nuclear contours. **G,** Lamin A/C intensity and nuclear envelope:nucleoplasm distribution ratios are elevated at BR compared to SR in HMC channels, whereas LMC nuclei show minimal regional differences. Intensity: HMC-BR, n = 17; HMC-SR, n = 18; LMC-BR, n = 20; LMC-SR, n = 20. Ratio: HMC-BR, n = 21; HMC-SR, n = 22; LMC-BR, n = 19; LMC-SR, n = 19. **H,** cPLA₂ accumulates at the nuclear envelope in BR nuclei under HMC conditions. **I,** Quantification reveals increased cPLA₂ peripheral distribution and higher nuclear Ca²⁺ intensity in BR nuclei, indicating activation of a Lamin A/C-cPLA₂-Ca²⁺ mechanotransductive loop. cPLA₂: HMC-BR, n = 23; HMC-SR, n = 21; LMC-BR, n = 16; LMC-SR, n = 16. Ca²⁺: HMC-BR, n = 20; HMC-SR, n = 20; LMC-BR, n = 18; LMC-SR, n = 17. Scale bars, 50 μm (A), 5 μm (F), 10 μm (H). Box plots show mean, quartiles and range; significance levels and non-significant results (ns) are indicated.

Under high-curvature (HMC) confinement, we observed that the nuclei become markedly distorted at the bend regions. Visually, the BR side of each nucleus appears indented or “scalloped” to conform to the curved channel wall, whereas the opposite side (SR) is less deformed (**Fig. 4B**). Quantitative shape analysis confirms that nuclei in HMC channels lose their circularity and develop high local curvature on the side facing the bend. In fact, the mean nuclear envelope curvature in BR segments (for HMC-confined cells) is significantly higher than that of SR segments in the same cells, and also higher than the nuclear curvature of any region in cells from low-curvature channels. 3D reconstructions of nuclei reveal alternating convex and concave curvature patches along the BR nuclear surface, indicating complex folding of the nuclear envelope in response to the undulating confinement (**Fig. 4C**). In contrast, nuclei in low-curvature (LMC) channels, as well as the SR portions of nuclei in HMC channels, remain much closer to spherical or ellipsoid shapes with minimal envelope bending.

To capture the overall effect, we measured a battery of nuclear morphometric parameters (18 features in total, including nuclear volume, surface area, aspect ratio, curvature variance, and bending energy). Using principal component analysis (PCA) on these features, we found that nuclei naturally cluster into two groups that correspond to their curvature context (**Fig. 4D**). One cluster (Cluster A) is defined by high nuclear bending energy, high curvature variance, and low sphericity, and this cluster consists predominantly (∼85%) of nuclei from the BR of HMC channels (**Fig. 4E**). The other cluster (Cluster B) comprises almost all nuclei from LMC channels and the SR portions of HMC nuclei, characterized by more moderate bending metrics and higher sphericity. In essence, HMC confinement pushes nuclei into a distinct morphological class, one could call them “high-curvature nuclei”, whereas LMC confinement allows nuclei to retain a baseline morphology. We calculated that bending energy in the nuclei experiencing HMC is over 30% higher than in LMC nuclei on average, and that nuclear shape irregularity (curvature variance) is strongly correlated with this bending energy and inversely correlated with nuclear roundness (**Supplementary Figs. S11-S12**). These data demonstrate that intense curvature confinement imposes a significant energetic and structural burden on the nucleus, enough to segregate nuclear morphologies into a separate category. Put simply, a nucleus can only bend so much before it must either initiate a mechanical response or risk damage; our results show that HMC conditions consistently drive nuclei into this extreme deformed state.

### Nuclear lamina reinforcement and mechanochemical response triggered by curvature stress

We next sought direct evidence of the nucleus activating mechanosignaling under curvature stress. Immunofluorescence imaging and live-cell reporters provided insight into how the nuclear envelope and associated signals responded in bend regions. In high-curvature channels, we found that Lamin A/C, the principal load-bearing component of the nuclear lamina, become locally enriched on the BR side of the nucleus, almost as if fortifying the envelope where it is most deformed (**Fig. 4F-G**). This Lamin-A/C accumulation coincides with spatial clustering of active mechanosignaling molecules. Specifically, cPLA₂ (cytosolic phospholipase A₂) shows localized recruitment to the nuclear envelope at BR sites, and we detected elevated nuclear Ca²⁺ levels in those same regions (using a fluorescent Ca²⁺ indicator) (**Fig. 4H-I, Supplementary Figs. S13A)**.

Notably, we did not observe a significant difference in YAP/TAZ (Yes-associated protein) localization between BR and SR or between HMC and LMC conditions (**Supplementary Figs. S13B**). YAP is a well-known mechanotransduction effector that shuttles to the nucleus in response to matrix stiffness or cell spreading, but in our curvature system YAP remains mostly nuclear in all conditions, and its activity does not appear to be specifically modulated by curvature. This suggests that the curvature-sensing pathway operates through a lamina-membrane tension-Ca²⁺ mechanism rather than through the YAP/TAZ pathway, highlighting a distinct axis of mechanotransduction engaged by geometric confinement.

In summary, high ECM curvature drives the nucleus into a state of maximal membrane tension and bending stress, which in turn activates a localized mechanochemical response. Even though cells in high-curvature channels move more slowly on a global scale, at the local scale their nuclei experience intense mechanical stimuli, essentially being repeatedly bent, and this triggers the nuclear envelope stretch response (Ca²⁺ release and cPLA₂ activation). In low-curvature channels, nuclei maintain their integrity and shape to a greater extent, avoiding the energy cost of continual deformation and thus allowing the cell to migrate efficiently without activating these stress pathways. This dichotomy illustrates how curvature can act as a highly localized mechanical cue: the overall cell might not feel “squeezed” in a large channel, but the nucleus can still detect the subtle curvature of its boundary and respond accordingly.

### Nuclear mechanosignaling under curvature reshapes cytoskeletal architecture and migration mechanics

Having established that high-curvature confinement induces pronounced nuclear bending and specific nuclear signaling events, we next examined how these nuclear mechanosignals feed back to the cell’s migration apparatus, namely, the cytoskeleton, focal adhesions, and traction forces that drive movement. We focused on comparing cells in high-curvature channels (where nuclear bending and mechanotransduction were evident) to those in low-curvature channels (where the nucleus remained less perturbed), reasoning those differences in the cytoskeletal architecture between these conditions would reveal how the nucleus’s sensing of curvature influences migration mechanics.

First, we looked at focal adhesion organization using paxillin staining, since paxillin is a key component of focal adhesions that was upregulated in the transcriptomic analysis. In low-curvature channels, paxillin-containing adhesions were distributed in small, discrete foci predominantly at the cell front and rear, typical of an elongated mesenchymal cell gripping the substrate (**Fig. 5A**). In high-curvature channels, however, we noticed that paxillin-rich adhesions became concentrated along the bend region of the cell, essentially forming continuous or belt-like adhesion structures aligned with the curved wall next to the bent nucleus. Quantification confirmed that paxillin intensity was ∼40-50% higher in the bend-region side of HMC cells compared to the side-region of those cells, whereas in LMC cells paxillin was evenly distributed with no significant regional difference (**Fig. 5B**). This indicates that focal adhesions are being spatially re-patterned by curvature: under high curvature, cells preferentially reinforce adhesions where the nucleus is experiencing stress, whereas under low curvature the adhesions remain more uniformly arranged to support straightforward locomotion.

**Figure 5.**
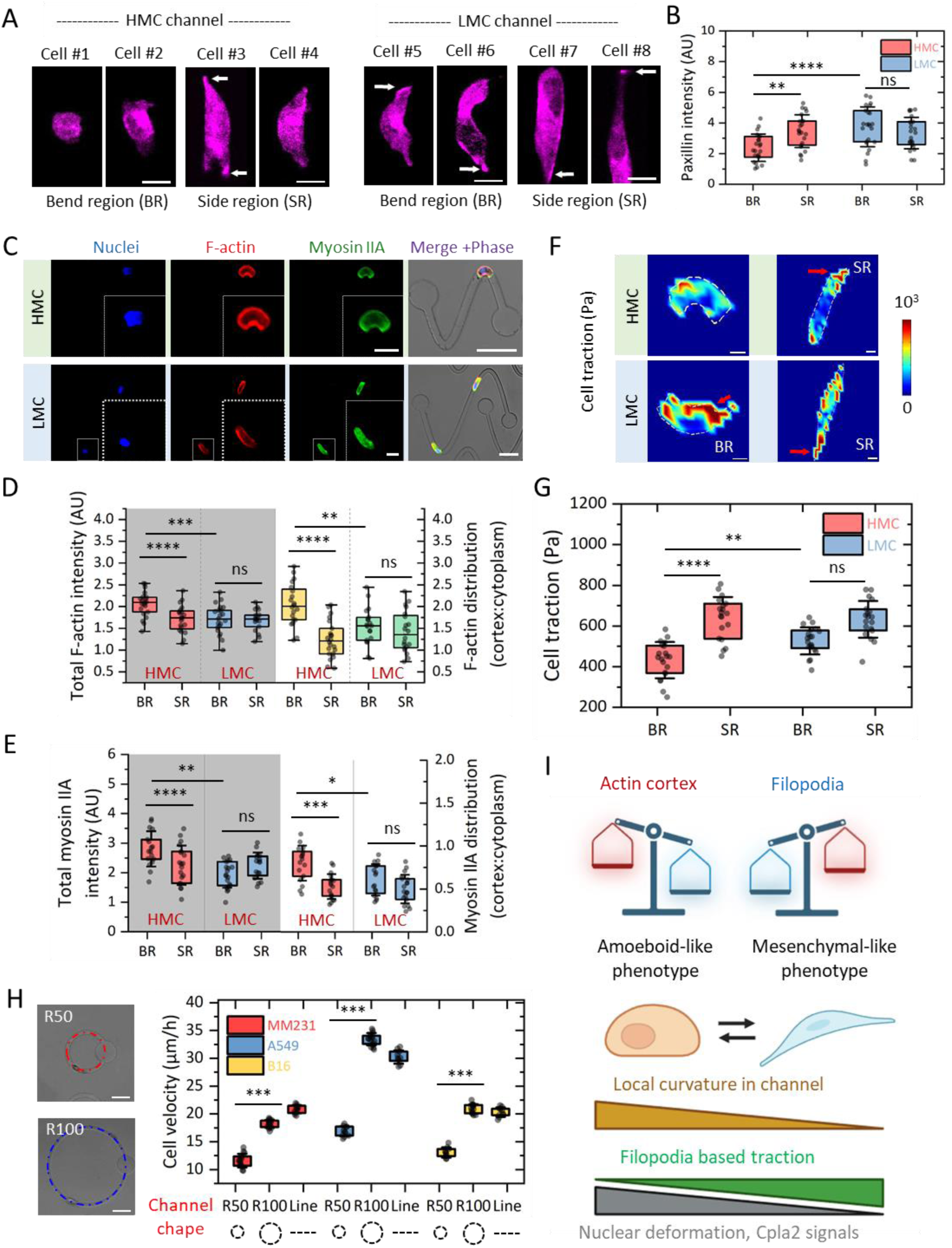
ECM curvature reprogrammes focal adhesions, actomyosin architecture and traction forces to switch migration phenotype. **A,** Paxillin immunofluorescence in cells migrating in high (HMC) and low (LMC) curvature channels reveals curvature-dependent adhesion patterns. In HMC channels, BR cells form dense paxillin belts aligned with the curved wall, whereas SR cells in the same channel and cells in LMC channels display more dispersed, axis-aligned foci. Arrows indicate focal adhesion enrichment. **B,** Paxillin intensity is ∼40–50% higher in BR(HMC) than SR(HMC), with no significant differences between BR and SR in LMC, showing that high curvature spatially biases adhesion reinforcement. HMC-BR, n = 23; HMC-SR, n = 23; LMC-BR, n = 22; LMC-SR, n = 22. **C,** Dual staining of F-actin (red) and myosin IIA (green), with nuclei (blue) and phase contrast, reveals distinct cytoskeletal architectures. HMC BR cells exhibit cortical actin bundles and myosin rings following the bent nuclear contour; LMC cells show prominent longitudinal stress fibres spanning the cell length. **D,** Total F-actin intensity and cortex:cytoplasm distribution ratios are elevated in BR(HMC) relative to SR(HMC), whereas LMC cells maintain lower cortical enrichment, indicating curvature-driven stabilisation of the actin cortex. Intensity: HMC-BR, n = 21; HMC-SR, n = 20; LMC-BR, n = 21; LMC-SR, n = 17. Ratio: HMC-BR, n = 20; HMC-SR, n = 22; LMC-BR, n = 19; LMC-SR, n = 20. **E,** Myosin IIA intensity and cortex:cytoplasm ratio are similarly increased in BR(HMC), supporting a shift toward cortex-dominated contractility in high curvature. Intensity: HMC-BR, n = 17; HMC-SR, n = 19; LMC-BR, n = 19; LMC-SR, n = 16. Ratio: HMC-BR, n = 17; HMC-SR, n = 18; LMC-BR, n = 16; LMC-SR, n = 19. **F,** Traction force microscopy maps show intense, spatially confined traction hotspots in BR(HMC) cells under paxillin belts, while SR(HMC) cells and LMC cells generate more diffuse or uniformly distributed traction patterns. **G,** Mean traction per cell is significantly higher in BR(HMC) than SR(HMC), whereas traction in LMC cells is more evenly distributed along the axis, consistent with persistent propulsion. HMC-BR, n = 20; HMC-SR, n = 19; LMC-BR, n = 20; LMC-SR, n = 19. **H,** Circular channels with defined radii (R50, high curvature; R100, low curvature) and linear tracks are used to generalise curvature effects. Across MDA-MB-231, A549 and B16F10 cells, migration velocity is markedly reduced in R50 compared to R100 and linear geometries, confirming that high curvature slows migration in diverse lineages. MM231 cells: R50, n = 25; R100, n = 25; Line, n = 21. A549 cells: R50, n = 21; R100, n = 21; Line, n = 21. B16F10 cells: R50, n = 32; R100, n = 32; Line, n = 32. **I,** Schematic model: low curvature favours elongated morphology, stress-fibre-based traction and mesenchymal-like persistent migration; high curvature promotes cortical actomyosin dominance, nuclear deformation and amoeboid-like exploratory movement. Scale bars, 20 μm (A, C), 10 μm (F), 50 μm (H). Box plots show median, quartiles and range; significance and ns values are indicated.

Next, we examined the actomyosin cytoskeleton. Low-curvature cells displayed prominent, stress fibres running longitudinally along the cell edges, connecting adhesion sites at the front and back (**Fig. 5C**). This architecture is ideal for generating strong, polarized traction forces for forward movement. In high-curvature cells, by contrast, stress fibres were less apparent; instead, we observed an increase in cortical actin, a shell of actomyosin around the cell periphery, particularly accentuated on the inner curvature side near the bent nucleus. Myosin-II (non-muscle myosin heavy chain IIA) localized robustly to this cortical network in HMC cells, consistent with the idea that nuclear signals were recruiting myosin to the cell cortex. In essence, high curvature confinement appears to shift the cell’s force-generation strategy from one dominated by filamentous stress-fiber contraction (in low curvature) to one dominated by cortical contraction (in high curvature). This shift was predicted by our mechanistic hypothesis: nuclear envelope stretch in HMC should activate RhoA and myosin via cPLA₂ and Ca²⁺, leading to increased cortical contractility. Our observations of the actin/myosin distribution support exactly this outcome (**Fig. 5D-E**).

We then measured the functional consequences on traction forces exerted by cells. Using traction force microscopy in the microchannels, we found that HMC BR cells generated strong, directional traction stresses at their front and rear ends, propelling them forward (**Fig. 5F**). However, SR(HMC) force maps showed weaker and more dispersed traction patches. In LMC cells, traction was relatively uniform along the axis in both BR and SR, with moderate intensity peaks supporting smooth, continuous propulsion. Mean traction (**Fig. 5G**) was significantly higher in BR(HMC) than SR(HMC), but total traction in LMC was more evenly distributed, aligning with the higher persistence seen in migration trajectories.

We next tested whether curvature alone could reprogram migration mechanics using circular microchannels with defined radii (R50 = high curvature, R100 = low curvature) versus linear tracks (**Fig. 5H**). Across MDA-MB-231, A549, and B16F10 cells, R50 channels markedly reduced migration velocity compared to R100 and linear geometries.

Together, these data indicate that ECM curvature governs not only nuclear mechanotransduction but also the spatial organisation of focal adhesions and actomyosin networks, which in turn define traction force patterns and migration efficiency. The schematic in **Fig. 5I** summarises these transitions: high curvature promotes cortical actin dominance, nuclear deformation, and amoeboid-like exploratory modes; low curvature favors filopodia-based traction, elongated cell morphology, and mesenchymal-like persistent migration.

### In vivo ECM curvature aligns nuclear orientation and induces bending

To assess whether curvature-dependent migration modes observed *in vitro* also manifest in native tissue, we examined the spatial relationship between nuclear morphology and extracellular matrix (collagen) architecture in orthotopic breast cancer tissue. Collagen fibres and nuclei were visualized in cryosections by immunofluorescence (**Fig. 6A-a**). Nuclear segmentation was performed using Cellpose (**Fig. 6A-b**), and ECM fibre orientation was extracted via our in-house curvature analysis pipeline (**Fig. 6A-c**). Overlay of nuclear orientation vectors with local ECM density revealed a strong alignment with most of nuclei from the nearest fibre direction (**Fig. 6A-d**).

**Figure 6.**
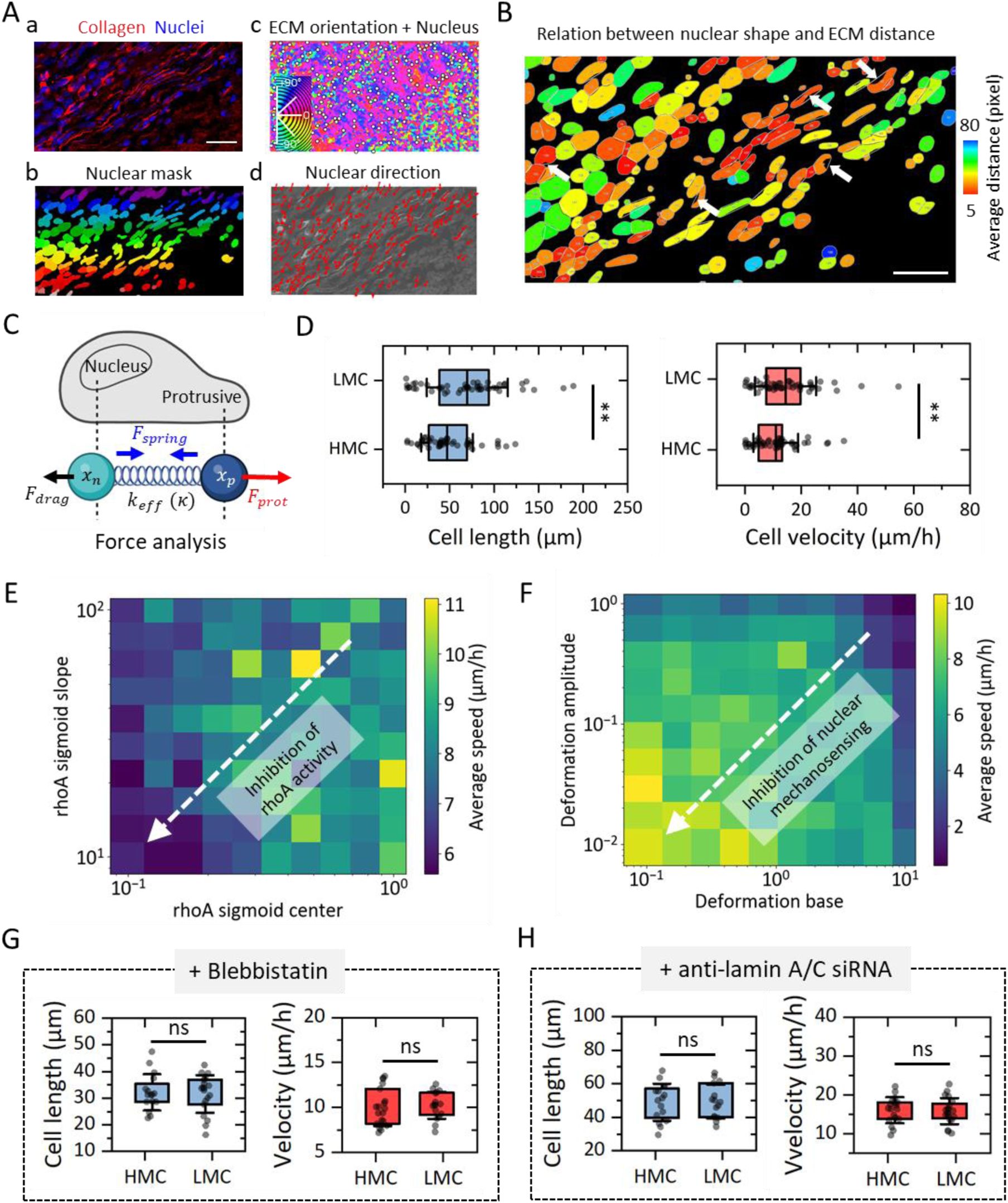
In vivo ECM curvature directs nuclear alignment and bending, and nuclear–cytoskeletal coupling controls curvature-dependent migration. **A,** In orthotopic breast tumours, immunofluorescence of ECM (collagen, red) and nuclei (DAPI, blue) reveals spatial relationships between fibre architecture and nuclear orientation. Cellpose-generated nuclear masks and principal nuclear orientation vectors overlaid on ECM orientation maps show preferential alignment of nuclei with local collagen fibres. **B,** Spatial mapping of the distance between nuclear contours and nearest ECM fibres (colour-coded from close, orange/red, to distant, blue) demonstrates that nuclei in direct contact with fibres are more elongated and bent, whereas distant nuclei remain rounded. **C,** A two-node active matter model represents nuclear and protrusive dynamics coupled by a curvature-dependent mechanotransduction term. The substrate is modelled as a sinusoidal trajectory whose curvature modulates RhoA activity, contractility and nuclear deformation costs. **D,** Simulations predict that cells on high-curvature substrates exhibit shorter length and reduced migration velocity compared to low-curvature substrates, mirroring experimental observations. Length: HMC, n = 50; LMC, n = 50. Velocity: n = 50; LMC, n = 50. **E-F,** Parameter sweeps show that migration speed depends non-monotonically on RhoA-mediated contractility and nuclear stiffness. Reducing curvature sensitivity or nuclear stiffness moves the system into a high-velocity regime, indicating that nuclear deformation cost is a primary energetic barrier to migration in complex topographies. **G-H,** Experimental perturbations validate the model. Pharmacological inhibition of ROCK (reducing actomyosin contractility) and siRNA-mediated Lamin A/C knockdown (softening the nucleus) both abolish curvature-linked differences in cell length and velocity between HMC and LMC channels. Add Blebbistatin: for length, HMC, n = 16; LMC, n = 19; for velocity: n = 19; LMC, n = 16. Add siRNA: for length, HMC, n = 16; LMC, n = 15; for velocity: n = 18; LMC, n = 20. Scale bars, 50 μm (A, B). Box plots depict median, quartiles and range; significance and ns values are indicated.

We next quantified the dependence of nuclear shape on proximity to ECM fibres. Population-wide morphometric analysis revealed a marked heterogeneity in nuclear shapes, with distributions skewed towards high eccentricity and aspect ratio, alongside a preferential orientation consistent with tissue architecture constraints (**Supplementary Fig. S14**). Spatially resolved mapping further elucidated the interaction between these shape parameters and ECM proximity (**Fig. 6B**). Nuclei located further from ECM structures were predominantly rounded, with low aspect ratios and minimal contour bending. In contrast, nuclei in close contact with ECM fibres displayed pronounced elongation and higher bending indices.

This observation was substantiated by feature correlation analysis, which demonstrated that deformation markers, such as mean invagination depth, were negatively correlated with the distance to the nearest ECM fibre, while geometric parameters like aspect ratio and solidity showed distinct dependencies on spatial positioning (**Supplementary Fig. S15**). These deformed nuclei often showed asymmetric curvature maps, with alternating convex and concave regions along the nuclear perimeter, closely resembling bent nuclear morphologies observed in vitro in confined microchannels. Together, these results confirm that ECM geometry *in vivo* directs nuclear alignment and deformation in a manner consistent with curvature-dependent migratory behaviour.

### Nuclear-cytoskeletal coupling mediates curvature-dependent cell migration in confinement

To elucidate the interplay between substrate curvature and cell migration in confined space, we implemented a stochastic two-node active matter model (**Fig. 6C**) where nuclear and protrusive dynamics are coupled via curvature-dependent mechanotransduction. Comparative simulations revealed that substrate curvature imposes significant physical constraints on cell migration (**Supplementary Fig. S16**). Cells migrating on HMC substrates exhibited a statistically significant reduction in both cell length and migration velocity compared to those on LMC substrates (**Fig. 6D**). This kinematic shift suggests that high local curvature triggers a “confinement response”, characterized by increased nuclear drag and cytoskeletal stiffening, which limits extension and retards forward progression.

We further explored the phase space of the model’s internal parameters to understand the robustness of this behavior. Varying the sensitivity of the RhoA signaling pathway to curvature (**Fig. 6E**) demonstrated that migration speed is non-monotonically dependent on contractility; the “inhibition of RhoA activity” trajectory highlights that shifting the system away from high-curvature-induced contractility can fundamentally alter the migration mode, though optimal velocity requires a precise balance between protrusion and retraction. Furthermore, the coupling between nuclear mechanics and geometry proved critical (**Fig. 6F**). Attenuating nuclear mechanosensing, specifically by reducing the deformation amplitude and base deformation parameters, effectively recovered migration speeds (indicated by the transition to the high-velocity yellow regime). This confirms that the physical cost of nuclear deformation acts as the primary energetic barrier to migration in complex topographies, a constraint that is alleviated when nuclear stiffness is uncoupled from geometric sensing.

To directly test the causal role of the nuclear mechanotransduction pathway in establishing these differences, we performed perturbation experiments. We used two complementary approaches: (*i*) reducing actomyosin contractility with a Rho-associated kinase inhibitor (ROCKi, which diminishes myosin-II activity), and (*ii*) knocking down Lamin A/C via siRNA to soften the nucleus and impair its mechanosensing ability (**Supplementary Fig. S17A**). Remarkably, both interventions eliminated the curvature-dependent differences in migration (**Fig. 6G-H**). ROCK inhibition caused highly contractile HMC-confined cells to become less contractile and behave more like they were in low curvature (**Supplementary Fig. S17B-E**): their speed increased and their directional persistence improved, essentially blunting the high-curvature-induced slowdown. Conversely, Lamin-A/C knockdown made the nucleus more deformable; cells with “softer” nuclei navigated high-curvature channels with less difficulty, maintaining higher speeds, and they no longer showed significant nuclear bending in curvature regions. These results confirm that the integrity of the nuclear-cytoskeletal coupling and contractile machinery is critical for curvature sensing. When the nucleus cannot properly sense or the cell cannot generate contractile response, the effect of curvature on migration is greatly diminished.

Taken together, our results support the idea that ECM curvature guides cell migration in tissues by imposing physical constraints on the nucleus (**Supplementary Fig. S18**). Regions of low curvature provide “fast lanes” where cells (including metastatic cancer cells or infiltrating immune cells) can move efficiently, aligning their nuclei and cytoskeleton for persistent migration. Regions of high curvature act as “obstacle courses” that force cells into a different migratory mode, one that might slow their advance but could aid in local exploration or tissue remodeling. Crucially, the entire curvature-sensing pathway depends on nuclear-cytoskeletal coupling: when that coupling is disrupted (experimentally or potentially by disease mutations), cells lose the ability to distinguish curvature differences, which could lead to maladaptive migration behavior.

## DISCUSSION

### Curvature sensing by the nucleus links stable tissue architecture to cellular migration strategies

Our study reveals a guidance cue for cell migration, *i.e.*, the geometric curvature of ECM fibres, and identifies the nucleus as the key mechanosensor that decodes this cue. We demonstrate that ECM curvature functions as a tissue-specific “geometric code” that remains remarkably stable even as tissues undergo pathological remodeling. This finding adds a new layer to the hierarchy of physical guidance cues in the microenvironment. Whereas cells frequently remodel ECM stiffness and alignment, fiber curvature appears to be an intrinsic property that cells encounter but do not easily erase. From a biophysical perspective, curvature is a structural feature set by how the tissue ECM is organized during development and constrained by anatomy, so it makes sense that cells have evolved a way to interpret this stable signal. By establishing that the nucleus can serve as a “curvature sensor”, our work unifies this concept with other forms of mechanotaxis. Previous research has shown that cells respond to matrix stiffness gradients (durotaxis) and topographic patterns (topotaxis/ratchetaxis), with the nucleus often playing a central role in these processes. We now place curvotaxis, directional migration governed by curvature gradients, into this framework and provide a mechanistic basis for it. A 2018 study had coined the term curvotaxis and observed that cells can preferentially migrate toward concave curvature areas on micropatterned surfaces^25^. Our findings substantially extend this concept by showing that in a 3D tissue-like context, nuclear mechanosensing underlies curvotactic behavior. In curved confinement, the nucleus effectively “feels” the degree of curvature by how much it must deform, and it transmits this information to the rest of the cell to dictate migration mode.

Mechanistically, our data support a model in which nuclear envelope stretching and tension are the proximate signal for curvature sensing. When a cell encounters a curved ECM constraint, the part of its nucleus abutting the curve experiences membrane unfolding and stress, analogous to what happens when a cell squeezes through a narrow pore^26–28^. This in turn triggers a cascade, Lamin-A/C reinforcement, Ca²⁺ release from the ER, and cPLA₂ recruitment to the inner nuclear membrane, ultimately leading to RhoA/myosin-II activation and cytoskeletal reorganization^29–30^. This pathway has been described in the context of cell confinement and compression by pioneering work in the field^24,28,31^, and our results now connect it to guidance by curvature. An intriguing aspect of this mechanism is that it operates independently of classical focal-adhesion-based or Hippo-pathway signaling. We found that YAP/TAZ, a well- known mechanotransduction effector sensitive to substrate stiffness and spread area, is not significantly involved in curvature sensing in our system. In soft or stiff 2D environments, YAP translocation to the nucleus correlates with cell mechanical state, but in our curved 3D channels, YAP was constitutively nuclear and did not distinguish high *vs.* low curvature. This suggests that the cell uses a specialized nuclear mechanosensory pathway (lamina-cPLA₂-Ca²⁺) for curvature, which might be evolutionarily tuned to detect confined geometry, whereas YAP/TAZ is tuned to substrate stiffness or 2D area. In essence, cells possess multiple mechanosensing “circuits”, and the nucleus-centric circuit comes to the forefront when navigating complex 3D topography.

From a physics standpoint, the nucleus acting as a curvature gauge aligns with the idea that it is the largest solid element inside a cell. It has a relatively fixed size and stiffness, so when external geometry is tight or highly curved, the nucleus accumulates bending energy (we measured >30% higher bending energy in high-curvature conditions). This stored energy and deformation then feedback as signals that alter cell mechanics. Our findings thus support a view of the nucleus as not just a genome container but also a physical ruler or rheostat that continuously measures the shape of the cell’s microenvironment. This principle of “nuclear curvotaxis” could be conserved across many cell types and tissues, explaining diverse observations such as why neutrophils slow down in highly tortuous tissue mazes or why cancer cells elongate in linear tracks but round up in irregular matrices. By coupling a stable geometric feature of tissues (curvature) to an adaptive cellular response (migration mode switching), nuclear curvotaxis provides a link between long-term tissue architecture and short-term cell behavior.

### Implications for cancer metastasis, immune infiltration, and tissue engineering

Our results carry several important implications for health and disease. In cancer biology, they suggest that the geometry of tumor ECM fibres could bias the routes and modes of cancer cell dissemination. For instance, a tumor with regions of low ECM curvature (long, aligned collagen fibres radiating outward) might effectively create “highways” that channel cancer cells into fast, persistent migration, potentially facilitating escape into blood vessels or lymphatics and thus promoting distant metastasis^32–33^. In contrast, a tumor microenvironment dominated by high-curvature matrices (*e.g.*, dense, looped fibres encircling tumor nests) might induce a more exploratory, amoeboid migration that confines cells to the local tissue or lymphatic niches. This could explain patterns where certain tumors preferentially spread locally versus systemically. It also resonates with clinical observations that collagen architecture correlates with patient outcomes: breast tumors with radially aligned, straight collagen fibres (low curvature) are associated with higher metastatic risk and poorer prognosis^34–37^, whereas highly disordered or curved collagen matrices can act as barriers to efficient dissemination^38–39^. Moreover, our finding that curvature can prime cells transcriptionally for invasiveness even while slowing them kinematically raises the possibility that high-curvature tumor regions might serve as “training grounds” where cancer cells acquire aggressive traits (like EMT and stemness) before moving to other areas.

In the context of immune cell infiltration, ECM curvature may similarly impact how immune cells navigate fibrotic or tumor tissues. Immune cells such as T cells often migrate along collagen fibres toward chemoattractant signals^40–42^. If those fibres are highly curved or form tangled networks, our results predict that T cells might adopt a less directed, slower migration, potentially hindering their ability to reach cancer cells. This is in line with the observation that dense, collagen-rich tumors tend to exhibit poor T-cell infiltration and are immunotherapeutically “cold”^43–45^. It is tempting to speculate that by modifying the ECM (for example, enzymatically digesting collagen crosslinks or re-aligning fibres), one might reduce curvature obstacles and enable immune cells to penetrate tumors more effectively. Our findings provide a mechanistic rationale for therapies aimed at remodeling tumor stroma: not only does ECM remodeling affect molecular signaling, but also the physical parameter of curvature could be optimized to either impede cancer cell migration or facilitate immune cell access.

From a broader tissue physiology perspective, developmental and regenerative processes may also leverage nuclear curvature sensing. During embryonic morphogenesis, cells migrate through geometrically complex environments (branching tubes, curved organ scaffolds)^46–49^. A conserved nuclear curvotaxis mechanism might help cells decide when to slow down and explore a niche versus when to move quickly to a target location. Likewise, in wound healing, fibroblasts encounter curved fibrin clots and provisional matrices; their behavior could be influenced by those geometries in ways that affect scar patterns^50–52^. It is noteworthy that we found tissue curvature fingerprints are set early in pathological remodeling, this could mean that the outcome of processes like fibrosis is imprinted by initial boundary conditions and continues to guide cell behavior throughout healing. Understanding this geometry-cell behavior coupling could open new avenues in tissue engineering: by designing biomaterial scaffolds with specific curvature patterns, we might direct cells to migrate in desired ways to promote organized tissue formation or prevent undesirable cell arrangements.

### A framework for predicting cell migration from ECM microstructure

A practical outcome of this work is the suggestion that we can predict how cells will migrate in a given tissue by quantifying its ECM curvature. Traditionally, mechanobiology has focused on stiffness or ligand density as predictors of cell movement, but those properties can be heterogenous and actively altered by cells. In contrast, measuring the curvature distribution of fibres in a tissue (for example, via imaging and computational skeleton analysis as we employed) yields a “curvature fingerprint” that is intrinsic to the tissue’s structure. Our data show that this fingerprint has functional significance: it governs whether cells in that tissue are more likely to migrate in a fast persistent mode or a slow exploratory mode. This insight could be developed into diagnostic or prognostic tools. For example, one could envision analyzing biopsies of tumor stroma to determine the mean fiber curvature; a tumor with an extremely low curvature ECM might be flagged as having a microenvironment conducive to metastasis (since it provides highways for cells to escape), whereas a high-curvature ECM tumor might have a different invasion pattern (more local or lymphatic spread). Such analysis could complement existing markers and perhaps guide treatment strategies (*e.g.*, more aggressive systemic therapy for tumors with “highway” ECM geometries). Similarly, in tissue engineering and implant design, incorporating the appropriate curvature cues could help guide cell migration and positioning. For instance, in designing a scaffold for nerve regeneration, one might include linear low-curvature conduits to encourage rapid axonal growth in a directed path, whereas for recreating a complex tissue interface, higher curvature features might promote local cell infiltration and matrix remodeling.

### Limitations and future directions

While our study illuminates a novel aspect of cell migration, it also has limitations that point to opportunities for further research. First, our set of analyzed tissues, though spanning multiple organs and pathologies, is not exhaustive. We focused on relatively static ECM configurations (like established tumors and scars). Tissues with highly dynamic ECM turnover, such as healing wounds, developing organs, or diseases like liver cirrhosis, might not maintain a single curvature fingerprint over time. It remains to be tested whether the curvature principle holds in environments where matrix is rapidly deposited and degraded. Future work could apply our curvature mapping methodology longitudinally in injury models or organoids to see if cells still respond to an average curvature in rapidly changing matrices, or if other cues dominate in those contexts. Broadening the range of tissues (including non-fibrillar matrices like brain perineuronal nets or basement membranes with inherent curvature in folds) will also reveal how universal nuclear curvotaxis is.

## RESOURCE AVAILABILITY

### Lead contact

Further information and requests for resources and reagents should be directed to and will be fulfilled by the lead contact, Feng Xu.

### Materials availability

All unique reagents generated in this study are available from the lead contact with a completed Materials Transfer Agreement.

### Data and code availability

- Single-cell RNA-seq data will be deposited at GEO once our paper is published..
- Original code for the ECM curvature analysis pipeline and the two-node active matter model will be deposited at GitHub once our paper is published.
- Any additional information required to reanalyze the data reported in this paper is available from the lead contact upon request.

## ACKNOWLEDGMENTS

The authors thank the patients who participated in this study and their families. This study was funded by the National Natural Science Foundation of China (no. 12572352 to B.C., no. 12225208 and no. 12432015 to F.X.), the Surface Project (Key Grant) of Chinese Medicine Education Assciation (no. 2024KTZ037) to Y.L., and The Youth Innovation Team of Shaanxi Universities, Technology Innovation Leading Program of Shaanxi (2024QCY-KXJ069) to L.W., Scientific Research Program Funded by Education Department of Shaanxi Provincial Government (24JP160) to L.W..

## AUTHOR CONTRIBUTIONS

B.C., Ye Li, Z.X., Y. Liu, L.W.K., Q.H.S., X.W., H.G., X.W., N.M.D., L.J.D., F.Y.L., C.Y.W., L.W., and Y.L. performed the experiments. B.C., Ye Li, Z.X., and Y. Liu analyzed the data. All autors reviewed and edited the original draft. M.L., T.W., T.J.L. and F.X. conceived the overall project, acquired funding, conceived the study, were responsible for administration and supervision.

## DECLARATION OF INTERESTS

The authors declare no competing interests.

## Supplemental Information

- Figures S1-S18 and Table S1.

## STAR★METHODS

Detailed methods are provided in the online version of this paper and include the following:

- **KEY RESOURCES TABLE**
- **EXPERIMENTAL MODELS AND STUDY PARTICIPANTS**

- Ethical statement
- Human subjects
- Animal models
- Cell lines and culture conditions
- **METHOD DETAILS**

- Fabrication of sinusoidal PEG microchannels
- Cell migration assay
- Immunofluorescence and confocal microscopy
- Ca²⁺ detection and quantitative analysis
- YAP nuclear localisation ratio analysis
- Pharmacological treatments
- siRNA-mediated knockdown of lamin A/C
- Traction force microscopy
- **QUANTIFICATION AND STATISTICAL ANALYSIS**

- Quantitative ECM curvature mapping
- Nuclear morphometry and curvature analysis
- Single-cell RNA Sequencing (scRNA-seq)
- Physical modeling of curvature-guided cell migration
- Statistics analysis

## STAR★METHODS

### KEY RESOURCES TABLE

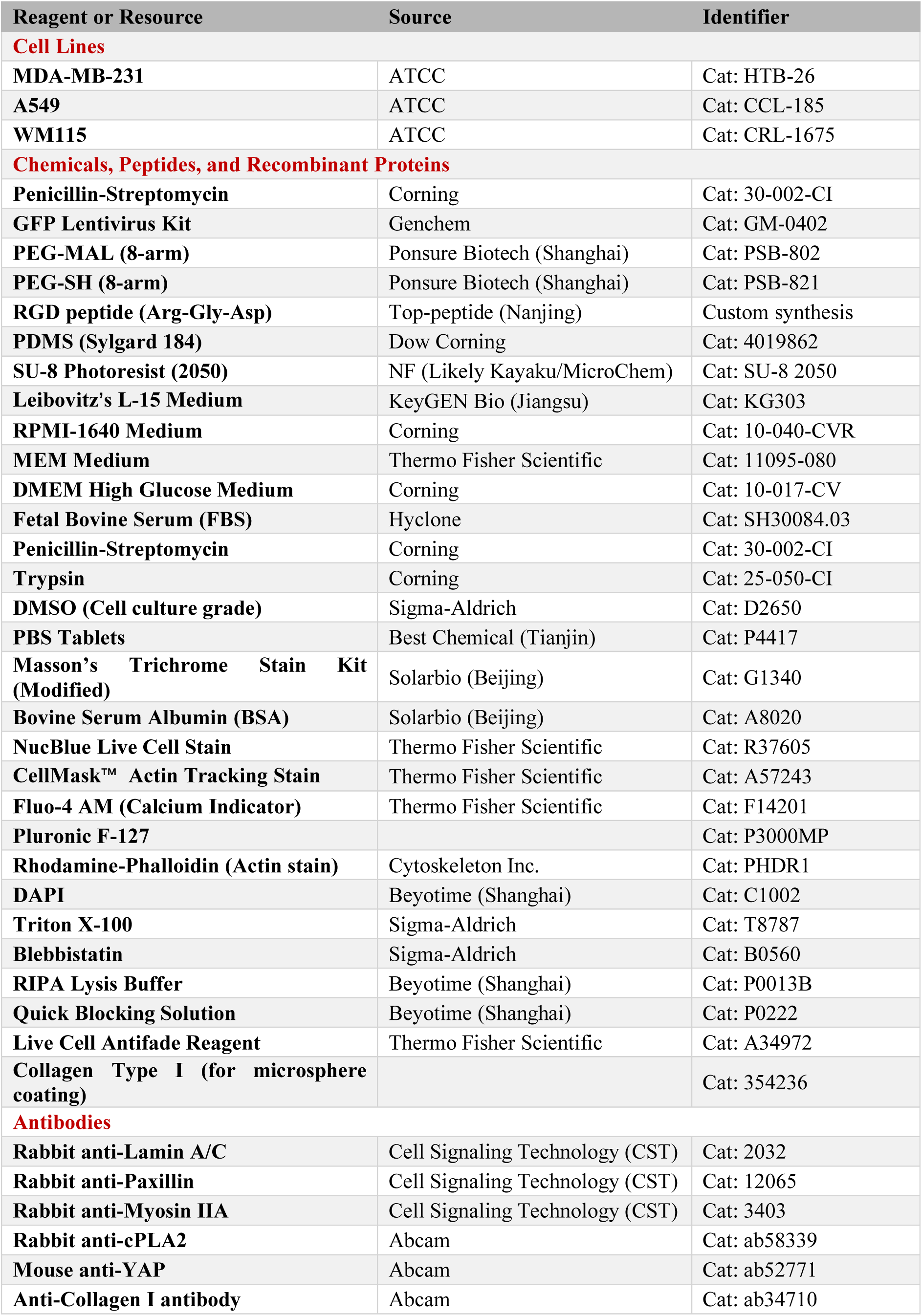
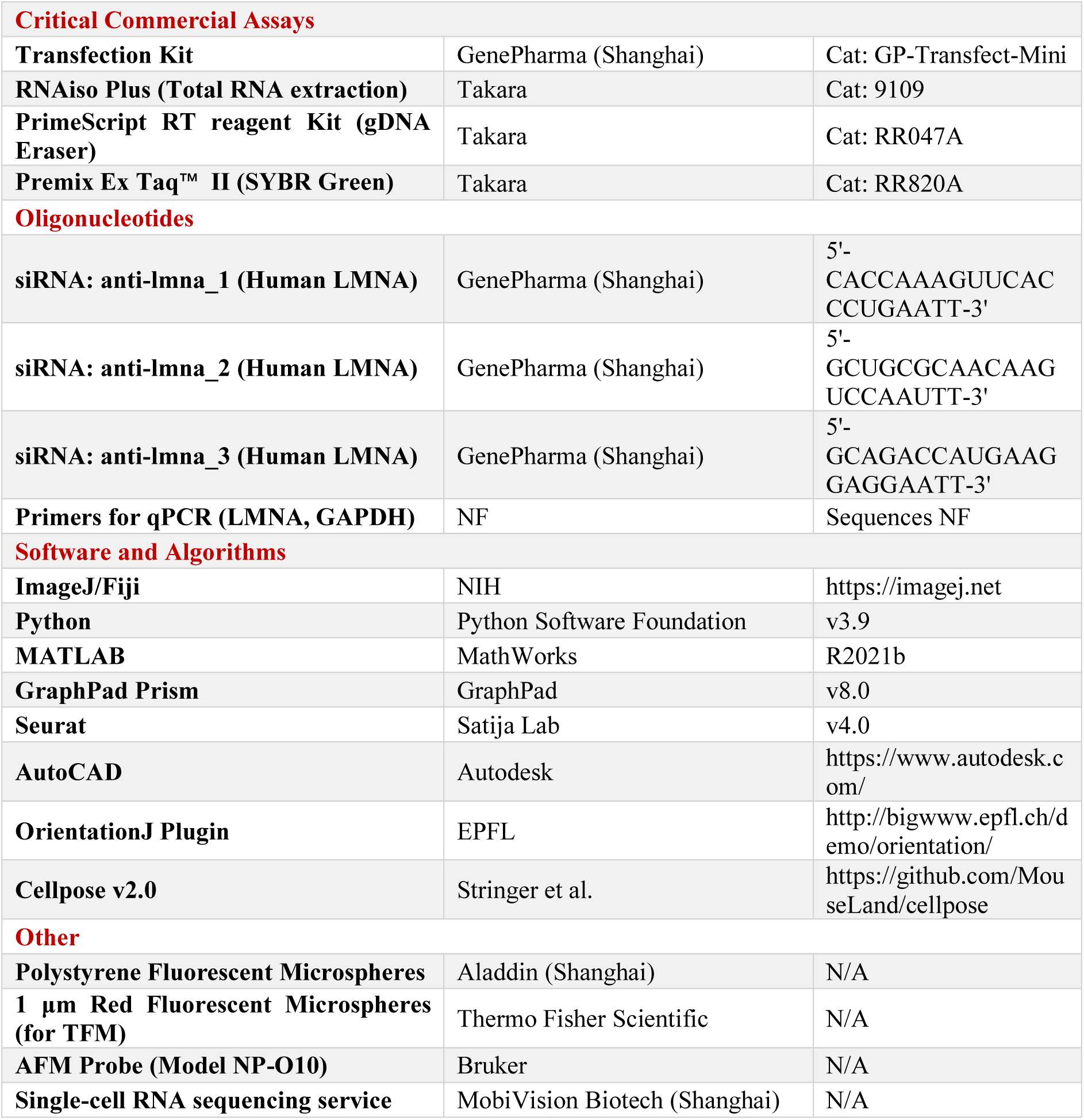

### EXPERIMENTAL MODEL AND STUDY PARTICIPANT DETAILS

#### Human subjects

Human tissue samples, including Triple-Negative Breast Cancer (TNBC), skin scar, gastric cancer, and lung puncture samples were collected from the First Affiliated Hospital of Xi’an Jiaotong University. Healthy gingival and periodontitis tissues were collected as controls. All samples were obtained with written informed consent and approved by the Biomedical Research Ethics Committee of School of Life Sciences and Technology in Xi’an Jiaotong University (Approval No. 2025-42). Samples were fixed in 4% paraformaldehyde (PFA) immediately after resection, embedded in paraffin, and sectioned at 4 µm for histological analysis.

#### Animal models

All animal experiments were performed in strict accordance with the guidelines of the Institutional Animal Care and Use Committee (IACUC) of Xi’an Jiaotong University (Protocol No. 2025-42).

- *<u>Melanoma Model</u>*: Female C57BL/6 mice (6-8 weeks old) were injected orthotopically (subcutaneously) with 1×10^6^ B16F10 cells (suspended in 50 µL PBS) into the dorsal skin. Tumors were surgically resected when diameters reached the ethical limit (>1.2 cm in preliminary observation, or strictly following IACUC humane endpoints).
- *<u>Breast Cancer Model</u>*: Female BALB/c nude mice (6-8 weeks old) were housed under SPF conditions. MDA-MB-231 cells (2×10^6^ cells in 50 µL PBS) were injected into the 4th inguinal mammary fat pad. Primary tumors were resected at the designated endpoint (∼8 weeks growth).

Post-operative care included analgesia and antibiotics according to veterinary guidelines. Lungs were harvested at study termination to assess metastasis via gross nodule counting and H&E staining.

#### Cell lines

- *<u>MDA-MB-231</u>*: Cultured in Leibovitz’s L-15 medium with 10% Fetal Bovine Serum (FBS, Hyclone) and 200 U/mL penicillin-streptomycin (Corning). Maintained at 37°C in free air exchange (no CO₂).
- *<u>A549</u>*: Cultured in RPMI-1640 medium with 10% FBS and 100 U/mL penicillin-streptomycin. Maintained at 37°C, 5% CO₂.
- *<u>B16F10</u>*: Cultured in MEM or DMEM with 10% FBS and 100 U/mL penicillin-streptomycin. Maintained at 37°C, 5% CO₂.

All cell lines were obtained from ATCC, authenticated by STR profiling, and tested negative for mycoplasma. For live-cell cytoplasmic labelling, GFP was stably expressed through lentiviral transduction using a GFP lentivirus kit (GeneChem, Shanghai, China) following the manufacturer’s instructions. Transduced cells were selected with puromycin (3 μg/ml for 72 h) and maintained thereafter with 1 μg/ml puromycin. For nuclear counterstaining in live cells, NucBlue Live Cell Stain (Thermo Fisher Scientific, USA) was added 8-12 h prior to imaging, incubated for 30 min at 37 °C.

### METHOD DETAILS

#### Fabrication of sinusoidal PEG microchannels

Silicon masters containing sinusoidal patterns (defined amplitude A and period T) and circular patterns were fabricated via standard photolithography using SU-8 2050 photoresist and chrome photomasks, then silanized with trichlorosilane vapor to facilitate demolding. PDMS (Sylgard 184) was mixed at a 10:1 ratio, degassed, poured over the master, cured at 70°C for 2 h, and cut to 14×14 mm stamps that were passivated with 2% BSA before use. For hydrogel molding, a precursor solution containing 2.5% (w/w) PEG-MAL (8-arm), 1.25% (w/w) PEG-SH (8-arm), and 10 mM RGD peptide (Top-peptide) was prepared in a glass-bottom dish, the PDMS stamp was brought into contact with the solution and leveled, and the assembly was incubated at room temperature for 2 h to allow thiol-maleimide crosslinking. After gently removing the stamps in sterile ethanol, separate flat PEG sheets were polymerized as top-covers, and channel dimensions (height, width, curvature) were verified by confocal microscopy (1 µm Z-increments) to ensure fidelity to the design.

#### Cell migration assay

PEG microchannels were sterilized in 75% ethanol for ≥2 h, rinsed three times with sterile PBS (5 min each), and pre-conditioned with complete culture medium at 37°C for ≥2 h to adsorb serum proteins. Cells were harvested and resuspended at 1×10⁶ cells/mL in complete medium, and 50 µL of suspension was pipetted onto the channel region. After keeping plates static for 10 min to allow sedimentation, pre-fabricated PEG hydrogel top-covers were aseptically placed to seal the channels, followed by gently adding 1 mL of complete medium along the dish wall. Migration assays commenced after 8 h of incubation, with imaging performed on an Olympus FV3000 confocal microscope equipped with a Tokai Hit stage-top incubator (37°C, 5% CO₂). Time-lapse images (DAPI channel for NucBlue-stained nuclei) were acquired using a 10× objective at 30-min intervals for 10 h. Nuclear centroids were tracked using the Manual Tracking plugin in Fiji, from which instantaneous velocity, mean speed, and Mean Squared Displacement (MSD) were calculated, and persistence time was derived from the velocity autocorrelation function.

#### Immunofluorescence and confocal microscopy

For staining, 10 h post-seeding, top-covers were removed and cells were fixed with freshly prepared 4% PFA in PBS for 10 min at room temperature, then permeabilized with 0.5% Triton X-100 in PBS for 15 min and blocked with rapid blocking solution (Beyotime) for 15 min. Primary antibodies (Anti-Lamin A/C, Anti-Paxillin, Anti-Myosin IIA, Anti-YAP, Anti-cPLA2) were diluted 1:200 and incubated overnight at 4°C, followed by incubation with Alexa Fluor-conjugated secondary antibodies (1:200) for 2 h at RT; nuclei were counterstained with DAPI (1:50,000) and F-actin with Rhodamine-Phalloidin (1:1000) for 1 h. To minimize optical distortion from the hydrogel, the glass bottom with hydrogel was detached and inverted onto a 100 µL PBS drop on a high-precision coverslip before acquiring Z-stacks using a Zeiss LSM880 (40×/1.3 NA oil objective) or Olympus FV3000.

#### Ca²⁺ detection and quantitative analysis

Cellular calcium dynamics during migration were monitored using the fluorescent indicator Fluo-4 AM. Cells were loaded with 5 µM Fluo-4 AM and 0.02% Pluronic F-127 in complete medium for 30 min at 37°C, followed by three washes with pre-warmed PBS. Live-cell Ca²⁺ imaging was conducted on an Olympus FV3000 confocal microscope. Nuclear and cytoplasmic regions of interest (ROIs) were segmented based on DAPI staining and whole-cell outlines, respectively. The average fluorescence intensity was measured for each ROI, and the nuclear calcium ratio was calculated as:

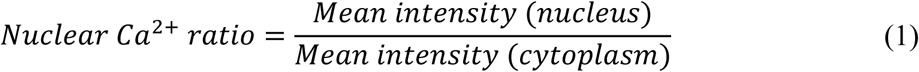

Background subtraction was performed for each channel using regions without cells. Image analysis was conducted in Fiji/ImageJ (NIH).

#### YAP nuclear localisation ratio analysis

Cells in microchannels were fixed and stained for YAP using anti-YAP antibody (, following the Immunofluorescence Staining protocol. Nuclei were counterstained with DAPI. Z-stack images were acquired by confocal microscopy and maximum-intensity projections generated. Nuclear ROIs were segmented from DAPI, and whole-cell ROIs from cytoplasmic fluorescence or phase-contrast boundaries. Mean intensity of YAP staining was measured within nuclear and total cell ROIs. The nuclear localisation ratio was determined by:

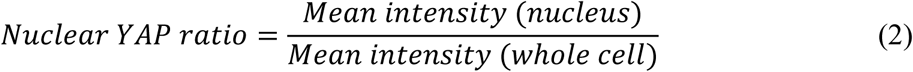

ROI boundaries were refined by morphological dilation/erosion and background-corrected.

#### Pharmacological treatments

To inhibit myosin II-mediated cortical contractility, cells were treated with 20 μM blebbistatin (Sigma-Aldrich) diluted in complete medium. Four hours before initiating live-cell tracking, culture medium in the PEG microchannel dishes was replaced with blebbistatin-containing medium following a PBS rinse. Inhibitor was maintained in the culture for the duration of imaging. Control cells were incubated with vehicle (DMSO, final concentration ≤0.1%).

#### siRNA-mediated knockdown of lamin A/C

Lamin A/C knockdown was performed using siRNA oligonucleotides designed against the human LMNA transcript (NM_170707). Three siRNA sequences were synthesized and screened, with one (siRNA3) selected for subsequent experiments due to its highest suppression efficiency. Cells at 30-40% confluence were transfected with 20 μM siRNA using a commercial transfection kit, following the manufacturer’s protocol. After 4-6 h, the transfection mixture was replaced with complete medium. Knockdown efficiency was quantified by RT-PCR 72 h post-transfection, normalized to GAPDH. For functional assays, cells were transfected 48 h prior to seeding into microchannels, and suppression was further verified by immunofluorescence.

#### Traction force microscopy

Red fluorescent beads (1 µm, Thermo Fisher) were embedded in the PEG hydrogel pre-polymer at a volume fraction of 0.1% (v/v) before crosslinking. Fluorescence images of beads and bright-field images of cells were first acquired in the stressed state, after which cells were lysed using 100 µL RIPA buffer for 2 min and beads were re-imaged at the exact same positions in the null/relaxed state. Bead displacement fields were calculated via Particle Image Velocimetry (PIV) in MATLAB, and traction stress maps were reconstructed by solving the inverse elastic problem using the storage modulus of the gel (confirmed by AFM indentation) as the elasticity parameter.

### QUANTIFICATION AND STATISTICAL ANALYSIS

#### Quantitative ECM curvature mapping

Masson’s trichrome stained slides were scanned using a whole-slide scanner, and color deconvolution was applied to separate the collagen channel (blue) from cellular components using optimized vectors. The collagen channel underwent Difference of Gaussians (DoG) filtering to enhance fiber structures, followed by adaptive thresholding to generate binary masks, while the TWOLBMI algorithm identified high-density matrix regions to exclude sparse background. The binary network was skeletonized, and local curvature (*κ*) was calculated for every pixel along the fiber skeleton as the inverse of the osculating circle radius, with a Gaussian smoothing window applied to reduce pixelation noise. For feature extraction, orientation was mapped using the OrientationJ plugin; geometric fingerprints including lacunarity, fiber endpoints, mean fiber thickness, and alignment index were computed to generate radar plots; and point-wise curvature values were compiled into histograms and fitted with Gaussian models to derive the mean curvature (*μ*) for each tissue type.

#### Nuclear morphometry and curvature analysis

##### In vitro analysis

3D nuclear surfaces were reconstructed from confocal Z-stacks using Python (OpenCV) and ImageJ to compute shape descriptors including area, aspect ratio, circularity (4π×Area/Perimeter²), and solidity; alignment was calculated as the cosine of the angle between the nuclear major axis and the channel axis; invagination metrics were derived by identifying concave segments through curvature sign changes, with defined as the Euclidean distance from the concavity apex to the convex hull; and the Lamin A/C ratio was defined as (mean intensity in the 1 µm peripheral rim) / (mean intensity in nucleoplasm).

##### In vivo analysis

1. ***Image preprocessing and segmentation***: Image analysis was performed using a custom Python pipeline (Python v3.x) integrating OpenCV, Scikit-image, and SciPy. Binary masks of nuclei were obtained using Cellpose (segmentation model) and imported as labeled 16- bit TIFF images. To remove segmentation artifacts, objects with an area smaller than 50 pixels or larger than 100,000 pixels were filtered out. The ECM channel (collagen) was preprocessed using a Gaussian filter (*σ* = 2.0) to reduce noise while preserving structural features.
2. ***Quantification of nuclear morphometry***: For each valid nucleus, geometric features were extracted using the “measure.regionprops” module from Scikit-image, encompassing basic shape descriptors (area, perimeter, major and minor axis length) and shape complexity measures: aspect ratio calculated as *MajorAxis*/*MinorAxis*; circularity defined as 4π×*Area*/*Perimeter*² (with 1 indicating a perfect circle); solidity representing the ratio of region area to convex hull area; and orientation as the angle between the nuclear major axis and the horizontal axis ranging from -90° to +90°.
3. ***ECM orientation and coherency analysis***: The local orientation of collagen fibres was computed using the Structure Tensor (Gradient-based) method, where structure tensor elements (*I_xx_*, *I_yy_*, *I_xy_*) were calculated from Sobel gradients of the Gaussian-smoothed ECM image within a local window (size = 32 pixels). Fibre orientation (*θ*_ECM_) was derived for each pixel using the formula *θ* = 0.5 arctan(2*I_xy_*/(*I_xx_* - *I_yy_*)), and coherency was calculated to measure local anisotropy (degree of alignment) of the fibres, ranging from 0 (isotropic) to 1 (highly aligned).
4. ***Nucleus-ECM interaction analysis*:** To quantify contact guidance and physical confinement, alignment analysis was performed by sampling the local ECM orientation within a 20×20 pixel window centered on each nucleus, defining an Alignment Score (S) based on the angular difference (Δ*θ*) between nuclear orientation and local ECM orientation as S = 1 - (Δ*θ*/90°), with scores ranging from 1 (perfect alignment) to 0 (perpendicular). Proximity analysis was conducted by generating a Euclidean distance map from the binarized ECM mask (thresholded at the 75th percentile of intensity) and computing the minimum, mean, and maximum distances from the nuclear boundary to the nearest ECM pixel.
5. ***Detection of nuclear envelope invaginations:*** To assess nuclear deformation induced by physical crowding, we analyzed convexity defects of the nuclear contour by generating a convex hull for each nucleus using “cv2.convexHull”, defining invaginations as concave deviations (defects) where the depth exceeded a threshold of 5 pixels, and recording metrics comprising the number of invaginations per nucleus and the *mean*/*maximum* invagination depth.
6. ***Spatial pattern statistics:*** The spatial distribution of nuclei was evaluated using the Nearest Neighbor Index (NNI), calculated as the ratio of the observed mean nearest neighbor distance to the expected mean distance for a random distribution (0.5/√𝜌, where ρ is the nuclear density), with values <1 indicating clustering and values >1 indicating dispersion.

#### Single-cell RNA Sequencing (scRNA-seq)

Polyacrylamide (PA) microspheres with diameters of 40 µm (R40) and 115 µm (R115) were fabricated using a flow-focusing microfluidic device and functionalized with Collagen I via sulfo-NHS activation under UV crosslinking. Cells were seeded on microspheres (1×10⁵ cells per 5×10⁴ beads), cultured for 12 h, and recovered by enzymatic detachment and filtration (40 µm strainers) to isolate bead-bound cells. For sequencing and analysis, libraries were prepared using the 10x Genomics Chromium platform and sequenced on Illumina NovaSeq; data were aligned to GRCh38 using Cell Ranger v6.1, and cells with 200<nFeature_RNA<7500 and mitochondrial genes <20% were retained for downstream analysis. Clustering was performed with Seurat v4.0 after LogNormalization and scaling, with dimensionality reduction via PCA and UMAP; pathway analysis used irGSEAwith the “ssgsea” method for Hallmark gene sets; WGCNA was performed on normalized data to identify curvature-correlated gene modules through soft-thresholding power fitting; and pseudotime differentiation scores were inferred using CytoTRACE (R).

#### Physical modeling of curvature-guided cell migration

##### 1. Model overview and assumptions

This simulation implements a stochastic, biophysical model of single-cell migration on a 1D sinusoidal substrate. The model couples substrate geometry (curvature) with intracellular biochemical signaling (Rac1/RhoA) and cytoskeletal mechanics to simulate topotaxis (migration guided by topographic features). Here are some key assumptions: (*i*) Overdamped dynamics: The cell operates in a low-Reynolds-number regime where inertia is negligible. Motion is determined by the balance of active protrusion forces, elastic restoring forces, and viscous drag. (*ii*) Two-node architecture: The cell is simplified into two mechanical nodes: the **Nucleus** (***x_n_***) and the **Protrusion** (***x_p_***), connected by a elastic spring. (*iii*) Local curvature sensing: The cell detects the local curvature of the substrate at the position of its nucleus. This curvature acts as the primary external cue regulating cell state. (*iv*) Mechanochemical feedback: Geometric cues (curvature) directly modulate biochemical states (Rac1 activity, RhoA activity, adhesion), which in turn alter mechanical parameters (force generation, stiffness, and drag).

##### 2. Geometric environment

The substrate is modeled as a sinusoidal wave defined by amplitude *A* and period *T*:

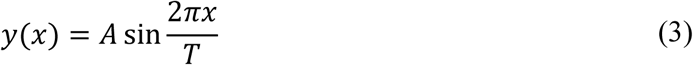

The local curvature κ(x) determines the geometric constraint felt by the cell. Analytically:

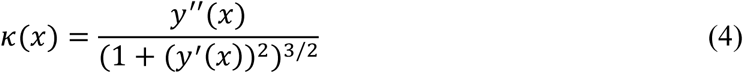

The model calculates this numerically using a finite difference method (central difference) to approximate the first (y′) and second (y′′) derivatives:

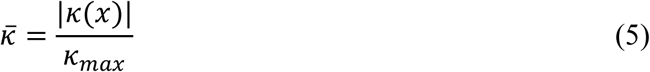

The curvature is normalized (𝜅̅) to a range [0, 1] based on a physiological threshold 𝜅𝑚𝑎𝑥 to drive the biochemical modules.

##### 3. Biomechanical modules (mechanotransduction)

We assume a competitive relationship between Rac1 (protrusion/polymerization) and RhoA (contraction/myosin II) in our model.

1. Rac1 activity (***R***): Inhibited by high curvature (simulating compressed actin networks in the protrusion).

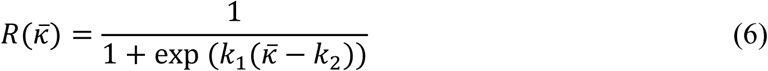
2. RhoA activity (***ρ***): Promoted by high curvature (simulating actin cortex formation).

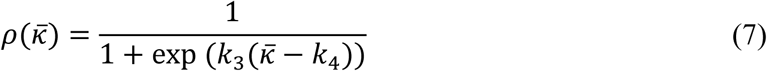
3. (3) Nuclear deformation (***δ*_def_**): Increases with curvature (steric compression).

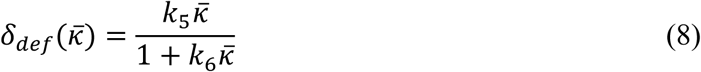

##### 4. Force balance and equations of motion

The motion of the cell is governed by the Langevin equation in the overdamped limit.

(1) Spring force (***F*_spring_**): Represents the cytoskeletal elasticity connecting the nucleus and protrusion. The stiffness is dynamic (Curvature Stiffening):

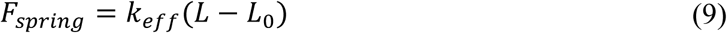

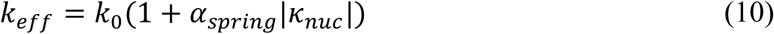

where *L*=*x_p_* - *x_n_* is current cell length. *L*_0_ is resting length. *α*_spring_ is sensitivity of stiffness to nuclear curvature.

(2) Protrusion force (***F_prot_***): Active driving force, generated only at the protrusion node (*x*_p_).

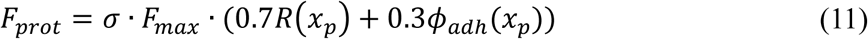

Where 𝜎 ∈ {−1,1} is the polarization direction. *F*_max_ is the maximum protrusion force capability.

(3) Nucleus drag: Variable, simulating steric hindrance (squeezing through narrow spaces).

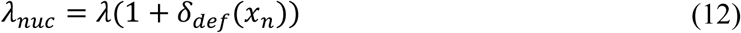

##### 5. System of equations

Applying Newton’s second law:

For the nucleus (*x_n_*),

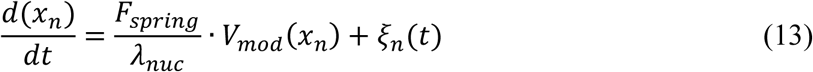

For the protrusion (*x_p_*),

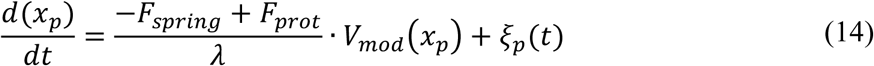

Where 𝜉(𝑡) represents Gaussian white noise modeling thermal fluctuations and intrinsic biological stochasticity:

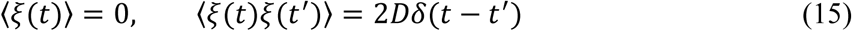

##### 6. Implementation details and output analysis

The simulation employs Euler-Maruyama integration with a time step of *d*t = 0.01 and a total of *N* = 5000 steps, outputting trajectories (*x*_n_(*t*), *x_p_*(*t*)), velocities, and instantaneous forces. Statistical significance is assessed using independent t-tests comparing “Small λ” (wavelength) versus “Large λ” conditions to evaluate the impact of topographical confinement on migration efficiency. Parameter table is in the supplementary materials.

#### Statistics analysis

Statistical analyses were performed using GraphPad Prism 8.0 and R (v4.1.2), with normality assessed using the Shapiro-Wilk test. For two-group comparisons, unpaired two-tailed Student’s t-test was used for normal data and Mann-Whitney U test for non-normal data, while multi-group comparisons employed one-way ANOVA with Tukey’s post-hoc test for normal data or Kruskal-Wallis test with Dunn’s post-hoc test for non-normal data. All data are presented as mean ± 1.5 SD and interquartile range (25% to 75%), with n≥3 independent biological replicates performed for all experiments. Significance was defined as *p<0.05, **p<0.01, ***p<0.001, ****p<0.0001.

## Notes

### Competing Interest Statement

The authors have declared no competing interest.

## REFERENCES

1. Hynes, R.O. (2009). The Extracellular Matrix: Not Just Pretty Fibrils. Science 326, 1216–1219. 10.1126/science.1176009.

2. Cukierman, E., Pankov, R., Stevens, D.R., and Yamada, K.M. (2001). Taking Cell-Matrix Adhesions to the Third Dimension. Science 294, 1708–1712. 10.1126/science.1064829.

3. Cheng, B., Li, M., Lin, M., Guo, H., and Xu, F. (2025). Mechanobiology across timescales. Nature Reviews Physics 7, 621–644. 10.1038/s42254-025-00874-w.

4. Sunyer, R., Conte, V., Escribano, J., Elosegui-Artola, A., Labernadie, A., Valon, L., Navajas, D., García-Aznar, J.M., Muñoz, J.J., Roca-Cusachs, P., and Trepat, X. (2016). Collective cell durotaxis emerges from long-range intercellular force transmission. Science 353, 1157–1161. doi:10.1126/science.aaf7119.

5. Isomursu, A., Park, K.-Y., Hou, J., Cheng, B., Mathieu, M., Shamsan, G.A., Fuller, B., Kasim, J., Mahmoodi, M.M., Lu, T.J., et al. (2022). Directed cell migration towards softer environments. Nature Materials 21, 1081–1090. 10.1038/s41563-022-01294-2.

6. Cheng, B., Wan, W., Huang, G., Li, Y., Genin, G.M., Mofrad, M.R.K., Lu, T.J., Xu, F., and Lin, M. Nanoscale integrin cluster dynamics controls cellular mechanosensing via FAKY397 phosphorylation. Science Advances 6, eaax1909. 10.1126/sciadv.aax1909.

7. Tabdanov, E.D., Puram, V.V., Win, Z., Alamgir, A., Alford, P.W., and Provenzano, P.P. (2018). Bimodal sensing of guidance cues in mechanically distinct microenvironments. Nature Communications 9, 4891. 10.1038/s41467-018-07290-y.

8. Charras, G., and Sahai, E. (2014). Physical influences of the extracellular environment on cell migration. Nature Reviews Molecular Cell Biology 15, 813–824. 10.1038/nrm3897.

9. Winkler, J., Abisoye-Ogunniyan, A., Metcalf, K.J., and Werb, Z. (2020). Concepts of extracellular matrix remodelling in tumour progression and metastasis. Nature Communications 11, 5120. 10.1038/s41467-020-18794-x.

10. Brett, E.A., Sauter, M.A., Machens, H.G., and Duscher, D. (2020). Tumor-associated collagen signatures: pushing tumor boundaries. Cancer Metab 8, 14. 10.1186/s40170-020-00221-w.

11. Maller, O., Drain, A.P., Barrett, A.S., Borgquist, S., Ruffell, B., Zakharevich, I., Pham, T.T., Gruosso, T., Kuasne, H., Lakins, J.N., et al. (2021). Tumour-associated macrophages drive stromal cell-dependent collagen crosslinking and stiffening to promote breast cancer aggression. Nature Materials 20, 548–559. 10.1038/s41563-020-00849-5.

12. Goetz, Jacky G., Minguet, S., Navarro-Lérida, I., Lazcano, Juan J., Samaniego, R., Calvo, E., Tello, M., Osteso-Ibáñez, T., Pellinen, T., Echarri, A., et al. (2011). Biomechanical Remodeling of the Microenvironment by Stromal Caveolin-1 Favors Tumor Invasion and Metastasis. Cell 146, 148–163. 10.1016/j.cell.2011.05.040.

13. Zhang, K., Corsa, C.A., Ponik, S.M., Prior, J.L., Piwnica-Worms, D., Eliceiri, K.W., Keely, P.J., and Longmore, G.D. (2013). The collagen receptor discoidin domain receptor 2 stabilizes SNAIL1 to facilitate breast cancer metastasis. Nature Cell Biology 15, 677–687. 10.1038/ncb2743.

14. Baptista, D., Teixeira, L., van Blitterswijk, C., Giselbrecht, S., and Truckenmüller, R. (2019). Overlooked? Underestimated? Effects of Substrate Curvature on Cell Behavior. Trends in Biotechnology 37, 838–854. 10.1016/j.tibtech.2019.01.006.

15. Messal, H.A., Alt, S., Ferreira, R.M.M., Gribben, C., Wang, V.M.-Y., Cotoi, C.G., Salbreux, G., and Behrens, A. (2019). Tissue curvature and apicobasal mechanical tension imbalance instruct cancer morphogenesis. Nature 566, 126–130. 10.1038/s41586-019-0891-2.

16. Kharaishvili, G., Simkova, D., Bouchalova, K., Gachechiladze, M., Narsia, N., and Bouchal, J. (2014). The role of cancer-associated fibroblasts, solid stress and other microenvironmental factors in tumor progression and therapy resistance. Cancer Cell Int 14, 41. 10.1186/1475-2867-14-41.

17. Fischer, T., Hayn, A., and Mierke, C.T. (2020). Effect of Nuclear Stiffness on Cell Mechanics and Migration of Human Breast Cancer Cells. Front Cell Dev Biol 8, 393. 10.3389/fcell.2020.00393.

18. Maiques, O., Sallan, M.C., Laddach, R., Pandya, P., Varela, A., Crosas-Molist, E., Barcelo, J., Courbot, O., Liu, Y., Graziani, V., et al. (2025). Matrix mechano-sensing at the invasive front induces a cytoskeletal and transcriptional memory supporting metastasis. Nature Communications 16, 1394. 10.1038/s41467-025-56299-7.

19. Fujimoto, H., Yoshihara, M., Rodgers, R., Iyoshi, S., Mogi, K., Miyamoto, E., Hayakawa, S., Hayashi, M., Nomura, S., Kitami, K., et al. (2024). Tumor-associated fibrosis: a unique mechanism promoting ovarian cancer metastasis and peritoneal dissemination. Cancer and Metastasis Reviews 43, 1037–1053. 10.1007/s10555-024-10169-8.

20. Devarasetty, M., Dominijanni, A., Herberg, S., Shelkey, E., Skardal, A., and Soker, S. (2020). Simulating the human colorectal cancer microenvironment in 3D tumor-stroma co-cultures in vitro and in vivo. Scientific Reports 10, 9832. 10.1038/s41598-020-66785-1.

21. Lomakin, A.J., Cattin, C.J., Cuvelier, D., Alraies, Z., Molina, M., Nader, G.P.F., Srivastava, N., Sáez, P.J., Garcia-Arcos, J.M., Zhitnyak, I.Y., et al. (2020). The nucleus acts as a ruler tailoring cell responses to spatial constraints. Science 370, eaba2894. 10.1126/science.aba2894.

22. Kalukula, Y., Stephens, A.D., Lammerding, J., and Gabriele, S. (2022). Mechanics and functional consequences of nuclear deformations. Nature Reviews Molecular Cell Biology 23, 583–602. 10.1038/s41580-022-00480-z.

23. Renkawitz, J., Kopf, A., Stopp, J., de Vries, I., Driscoll, M.K., Merrin, J., Hauschild, R., Welf, E.S., Danuser, G., Fiolka, R., and Sixt, M. (2019). Nuclear positioning facilitates amoeboid migration along the path of least resistance. Nature 568, 546–550. 10.1038/s41586-019-1087-5.

24. Venturini, V., Pezzano, F., Català Castro, F., Häkkinen, H.-M., Jiménez-Delgado, S., Colomer-Rosell, M., Marro, M., Tolosa-Ramon, Q., Paz-López, S., Valverde, M.A., et al. (2020). The nucleus measures shape changes for cellular proprioception to control dynamic cell behavior. Science 370, eaba2644. 10.1126/science.aba2644.

25. Vassaux, M., Pieuchot, L., Anselme, K., Bigerelle, M., and Milan, J.L. (2019). A Biophysical Model for Curvature-Guided Cell Migration. Biophys J 117, 1136–1144. 10.1016/j.bpj.2019.07.022.

26. Infante, E., Castagnino, A., Ferrari, R., Monteiro, P., Agüera-González, S., Paul-Gilloteaux, P., Domingues, M.J., Maiuri, P., Raab, M., Shanahan, C.M., et al. (2018). LINC complex-Lis1 interplay controls MT1-MMP matrix digest-on-demand response for confined tumor cell migration. Nature Communications 9, 2443. 10.1038/s41467-018-04865-7.

27. Pieuchot, L., Marteau, J., Guignandon, A., Dos Santos, T., Brigaud, I., Chauvy, P.-F., Cloatre, T., Ponche, A., Petithory, T., Rougerie, P., et al. (2018). Curvotaxis directs cell migration through cell-scale curvature landscapes. Nature Communications 9, 3995. 10.1038/s41467-018-06494-6.

28. Callan-Jones, A.C., and Voituriez, R. (2016). Actin flows in cell migration: from locomotion and polarity to trajectories. Current Opinion in Cell Biology 38, 12–17. 10.1016/j.ceb.2016.01.003.

29. Provenzano, P.P., Inman, D.R., Eliceiri, K.W., Trier, S.M., and Keely, P.J. (2008). Contact guidance mediated three-dimensional cell migration is regulated by Rho/ROCK-dependent matrix reorganization. Biophys J 95, 5374–5384. 10.1529/biophysj.108.133116.

30. Swift, J., Ivanovska, I.L., Buxboim, A., Harada, T., Dingal, P.C.D.P., Pinter, J., Pajerowski, J.D., Spinler, K.R., Shin, J.-W., Tewari, M., et al. (2013). Nuclear Lamin-A Scales with Tissue Stiffness and Enhances Matrix-Directed Differentiation. Science 341, 1240104. 10.1126/science.1240104.

31. Reversat, A., Gaertner, F., Merrin, J., Stopp, J., Tasciyan, S., Aguilera, J., de Vries, I., Hauschild, R., Hons, M., Piel, M., et al. (2020). Cellular locomotion using environmental topography. Nature 582, 582–585. 10.1038/s41586-020-2283-z.

32. Weigelin, B., Bakker, G.J., and Friedl, P. (2012). Intravital third harmonic generation microscopy of collective melanoma cell invasion: Principles of interface guidance and microvesicle dynamics. Intravital 1, 32–43. 10.4161/intv.21223.

33. Lee, M., Downes, A., Chau, Y.Y., Serrels, B., Hastie, N., Elfick, A., Brunton, V., Frame, M., and Serrels, A. (2015). In vivo imaging of the tumor and its associated microenvironment using combined CARS / 2-photon microscopy. Intravital 4, e1055430. 10.1080/21659087.2015.1055430.

34. Heydari, S., Tajik, F., Safaei, S., Kamani, F., Karami, B., Dorafshan, S., Madjd, Z., and Ghods, R. (2025). The association between tumor-stromal collagen features and the clinical outcomes of patients with breast cancer: a systematic review. Breast Cancer Res 27, 69. 10.1186/s13058-025-02017-6.

35. Jiang, H., Hegde, S., Knolhoff, B.L., Zhu, Y., Herndon, J.M., Meyer, M.A., Nywening, T.M., Hawkins, W.G., Shapiro, I.M., Weaver, D.T., et al. (2016). Targeting focal adhesion kinase renders pancreatic cancers responsive to checkpoint immunotherapy. Nature Medicine 22, 851–860. 10.1038/nm.4123.

36. Joyce, J.A., and Fearon, D.T. (2015). T cell exclusion, immune privilege, and the tumor microenvironment. Science 348, 74–80. 10.1126/science.aaa6204.

37. Binnewies, M., Roberts, E.W., Kersten, K., Chan, V., Fearon, D.F., Merad, M., Coussens, L.M., Gabrilovich, D.I., Ostrand-Rosenberg, S., Hedrick, C.C., et al. (2018). Understanding the tumor immune microenvironment (TIME) for effective therapy. Nature Medicine 24, 541–550. 10.1038/s41591-018-0014-x.

38. Levental, K.R., Yu, H., Kass, L., Lakins, J.N., Egeblad, M., Erler, J.T., Fong, S.F.T., Csiszar, K., Giaccia, A., Weninger, W., et al. (2009). Matrix Crosslinking Forces Tumor Progression by Enhancing Integrin Signaling. Cell 139, 891–906. 10.1016/j.cell.2009.10.027.

39. Cui, G., Deng, S., Zhang, B., Wang, M., Lin, Z., Lan, X., Li, Z., Yao, G., Yu, M., and Yan, J. (2024). Overcoming the Tumor Collagen Barriers: A Multistage Drug Delivery Strategy for DDR1-Mediated Resistant Colorectal Cancer Therapy. Advanced Science 11, 2402107. 10.1002/advs.202402107.

40. Friedl, P., and Weigelin, B. (2008). Interstitial leukocyte migration and immune function. Nature Immunology 9, 960–969. 10.1038/ni.f.212.

41. Hartmann, N., Giese, N.A., Giese, T., Poschke, I., Offringa, R., Werner, J., and Ryschich, E. (2014). Prevailing Role of Contact Guidance in Intrastromal T-cell Trapping in Human Pancreatic Cancer. Clinical Cancer Research 20, 3422–3433. 10.1158/1078-0432.CCR-13-2972.

42. Tabdanov, E.D., Rodríguez-Merced, N.J., Cartagena-Rivera, A.X., Puram, V.V., Callaway, M.K., Ensminger, E.A., Pomeroy, E.J., Yamamoto, K., Lahr, W.S., Webber, B.R., et al. (2021). Engineering T cells to enhance 3D migration through structurally and mechanically complex tumor microenvironments. Nature Communications 12, 2815. 10.1038/s41467-021-22985-5.

43. Wang, S., Li, J., and Zhao, Y. (2025). Targeting collagen in “armored and cold” tumors: Overcoming barriers to cancer therapy. Cancer Pathogenesis and Therapy 3, 383–391. 10.1016/j.cpt.2024.11.001.

44. Peng, D.H., Rodriguez, B.L., Diao, L., Chen, L., Wang, J., Byers, L.A., Wei, Y., Chapman, H.A., Yamauchi, M., Behrens, C., et al. (2020). Collagen promotes anti-PD-1/PD-L1 resistance in cancer through LAIR1-dependent CD8+ T cell exhaustion. Nature Communications 11, 4520. 10.1038/s41467-020-18298-8.

45. Sun, X., Wu, B., Chiang, H.-C., Deng, H., Zhang, X., Xiong, W., Liu, J., Rozeboom, A.M., Harris, B.T., Blommaert, E., et al. (2021). Tumour DDR1 promotes collagen fibre alignment to instigate immune exclusion. Nature 599, 673–678. 10.1038/s41586-021-04057-2.

46. Ventura, G., and Sedzinski, J. (2022). Emerging concepts on the mechanical interplay between migrating cells and microenvironment in vivo. Front Cell Dev Biol 10, 961460. 10.3389/fcell.2022.961460.

47. Ishii, M., Tateya, T., Matsuda, M., and Hirashima, T. (2021). Stalling interkinetic nuclear migration in curved pseudostratified epithelium of developing cochlea. Royal Society Open Science 8, 211024. 10.1098/rsos.211024.

48. Luciano, M., Versaevel, M., Kalukula, Y., and Gabriele, S. (2024). Mechanoresponse of Curved Epithelial Monolayers Lining Bowl-Shaped 3D Microwells. Advanced Healthcare Materials 13, 2203377. 10.1002/adhm.202203377.

49. Yang, Y., Xu, T., Bei, H.-P., Zhang, L., Tang, C.-Y., Zhang, M., Xu, C., Bian, L., Yeung, K.W.-K., Fuh, J.Y.H., and Zhao, X. (2022). Gaussian curvature–driven direction of cell fate toward osteogenesis with triply periodic minimal surface scaffolds. Proceedings of the National Academy of Sciences 119, e2206684119. 10.1073/pnas.2206684119.

50. Xu, H., Huo, Y., Zhou, Q., Wang, L.A., Cai, P., Doss, B., Huang, C., and Hsia, K.J. (2023). Geometry-mediated bridging drives nonadhesive stripe wound healing. Proceedings of the National Academy of Sciences 120, e2221040120. 10.1073/pnas.2221040120.

51. Kollmannsberger, P., Bidan, C.M., Dunlop, J.W.C., Fratzl, P., and Vogel, V. Tensile forces drive a reversible fibroblast-to-myofibroblast transition during tissue growth in engineered clefts. Science Advances 4, eaao4881. 10.1126/sciadv.aao4881.

52. Wang, Z., Lauko, J., Kijas, A.W., Gilbert, E.P., Turunen, P., Yegappan, R., Zou, D., Mata, J., and Rowan, A.E. (2023). Snake venom-defined fibrin architecture dictates fibroblast survival and differentiation. Nature Communications 14, 1029. 10.1038/s41467-023-36437-9.

