## Supplementary Materials (PDF) for "A Conserved Geometric Code: Extracellular Matrix Curvature Directs Cell Migration Strategy via Nuclear Mechanosensing"

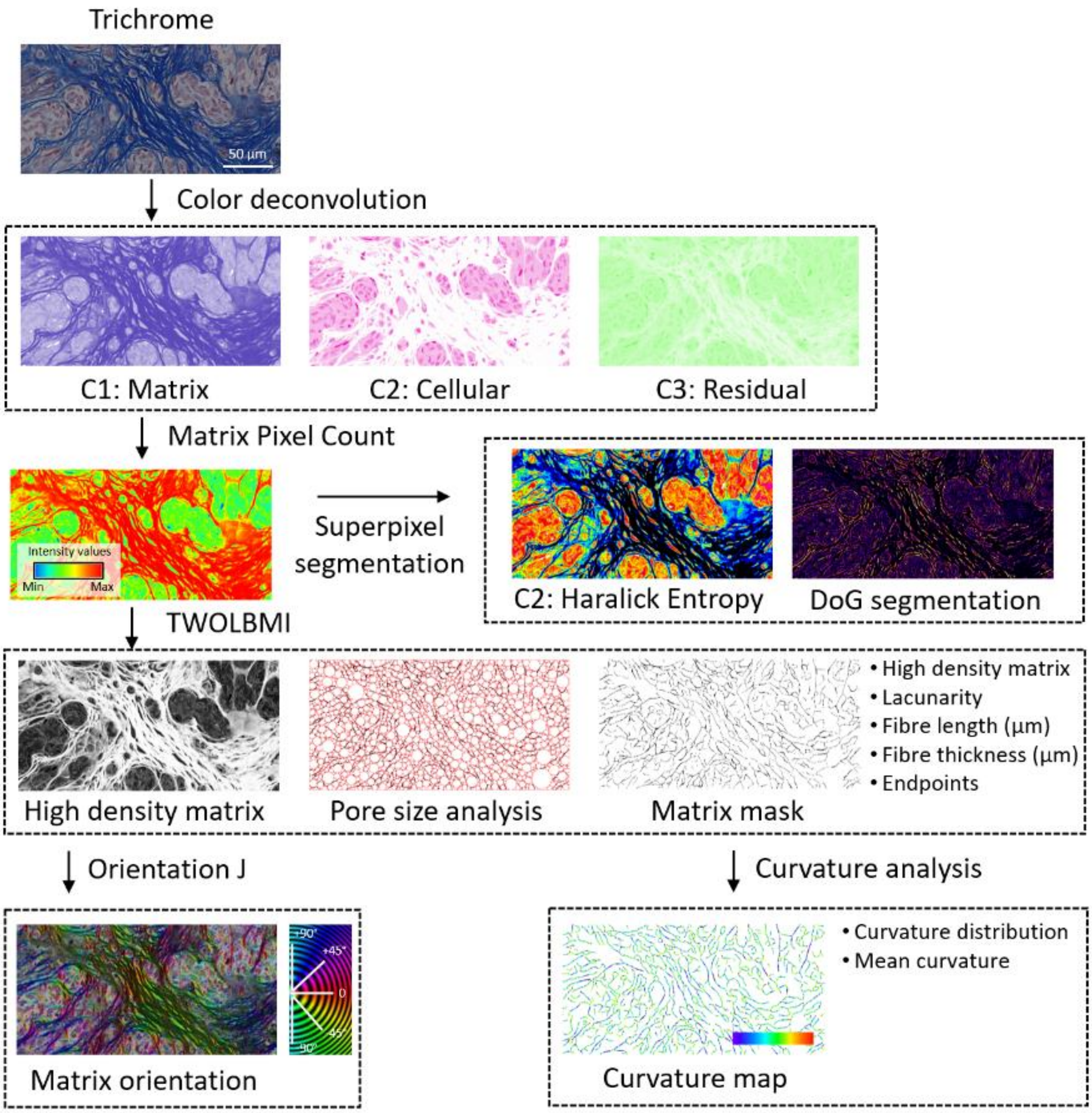


**Supplementary Fig. S1. Representative pipeline applied to tumour ECM sections stained with Masson's trichrome.** Color deconvolution separates matrix (C1), cellular (C2), and residual components (C3). Matrix pixel counting and superpixel segmentation are used to compute texture features (Haralick entropy) and DoG-based fibre segmentation. High-density matrix regions are identified via TWOLBMI, followed by pore-size analysis and extraction of a binary matrix mask. Fibre metrics (length, thickness, endpoints, lacunarity) are calculated, and OrientationJ assigns local fiber orientations (color-coded angle map). Curvature analysis generates pixel-wise curvature maps and summarizes curvature distributions, reporting mean curvature for each region.

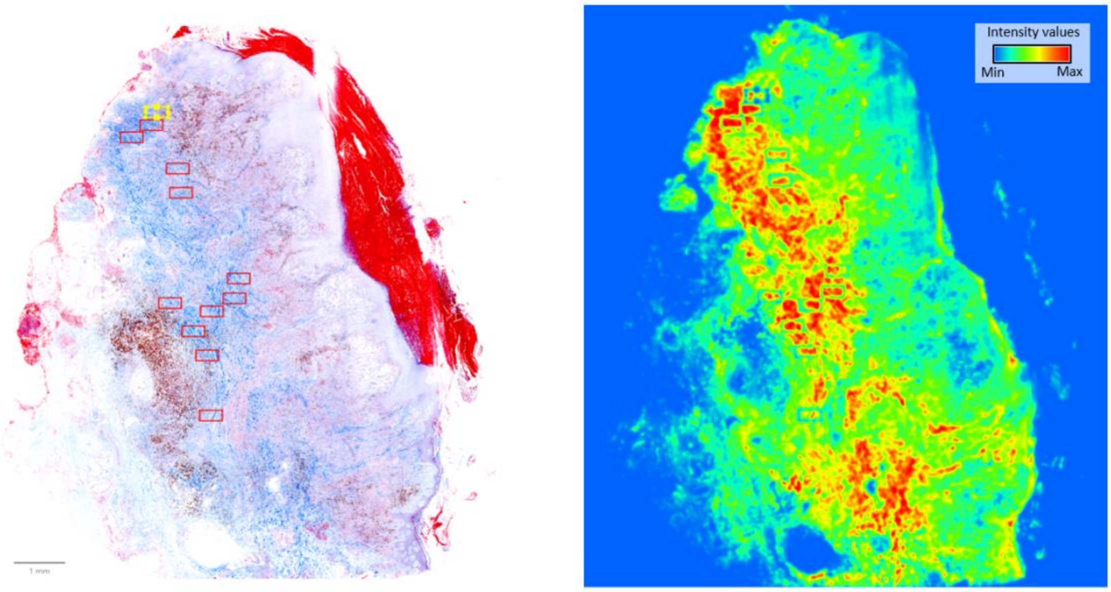


**Supplementary Fig. S2. Matrix region selection for ECM curvature analysis.** Representative whole-section view of Masson's trichrome-stained tumour tissue used for extracellular matrix (ECM) mapping. Left: Manually and semi-automatically defined rectangular regions (red boxes) mark areas of interest for curvature and texture profiling; selection was guided by histological ECM distribution and exclusion of artefacts. Right: Corresponding intensity map of the matrix channel (C1, post-color-deconvolution) showing pixel intensity values from minimum (blue) to maximum (red), highlighting zones of high-density ECM suitable for further fibre segmentation, orientation analysis, and curvature measurement (Methods). Scale bar: 1 mm (left panel). Intensity values are dimensionless (normalized grey-scale range).

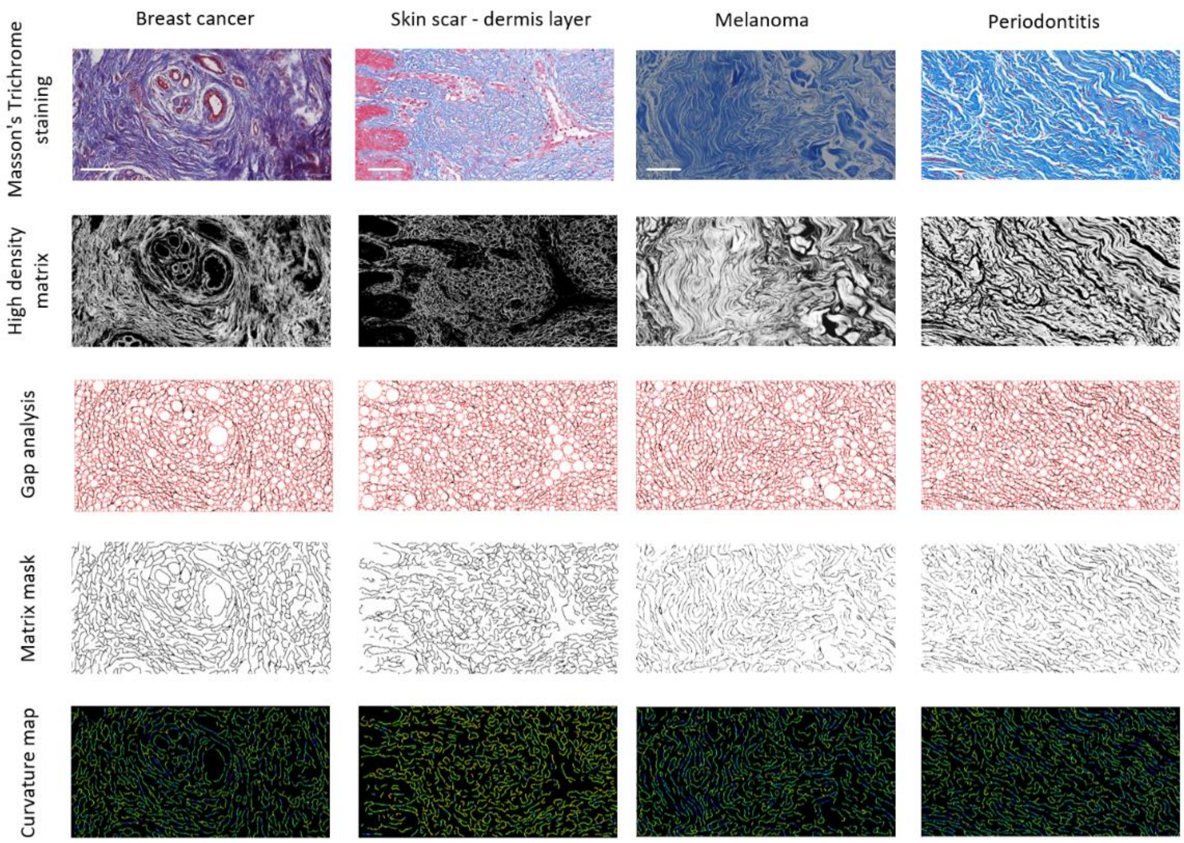


**Supplementary Fig. S3. Comparative ECM architecture and curvature profiles across diverse pathological tissues.** Representative extracellular matrix (ECM) analyses from histological sections of breast cancer, skin scar (dermis), melanoma, and periodontitis, processed using the standard pipeline (color deconvolution, high-density matrix extraction, gap analysis, mask generation, and curvature mapping; see Methods). Row 1: Masson's trichrome staining highlighting collagen-rich ECM (blue) and cellular regions (red). Row 2: High-density matrix maps showing compact fibre networks in black on a white background, indicating regions selected for subsequent morphometric analysis. Row 3: Gap analysis visualizing pore-size distribution (red outlines) within the ECM, where larger gaps correspond to lower fibre density. Row 4: Binary matrix masks derived from segmentation, delineating the continuous ECM scaffold from cellular spaces. Row 5: Curvature maps of ECM fibers, with color-coded local mean curvature (green to yellow; increasing curvature magnitude), revealing tissue-specific geometric differences in fiber organization. These comparative maps illustrate that ECM density, pore size, and fibre curvature vary substantially between tissue types and pathological states, potentially influencing in-situ nuclear morphology, cellular mechanosensing, and migration dynamics. Scale bars: 50 μm (all histological and analysis panels).

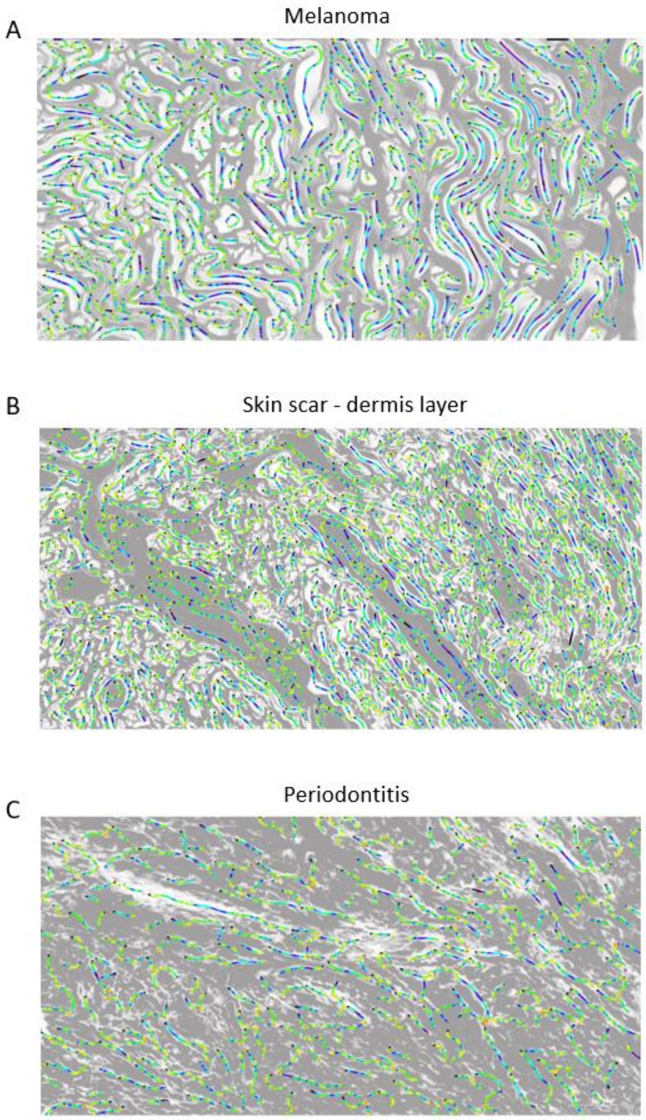


**Supplementary Fig. S4. Orientation mapping of ECM fibres in pathological tissue sections**. Representative orientation maps of collagen-rich extracellular matrix (ECM) fibres generated using the OrientationJ plugin from Masson's trichrome-stained histological sections. (**A**) Melanoma: Dense, highly curved fibre bundles with mixed orientation angles, color-coded according to the local principal direction (green to blue scale). (**B**) Skin scar (dermis): Heterogeneous fibre alignment with both tightly curved clusters and extended linear elements; indicative of scar remodelling. (**C**) Periodontitis: Predominantly elongated fibre structures with lower curvature variability, reflecting chronic inflammatory ECM reorganization. In all cases, grey underlay shows the high-density matrix mask, and color overlays depict the orientation angle of each fibre segment. These maps provide direct visual confirmation of ECM geometric diversity between tissues, reinforcing the in vivo findings in Fig. 6 that nuclear orientation aligns with local ECM fibre direction and curvature. Scale bars: 50 µm (all panels).

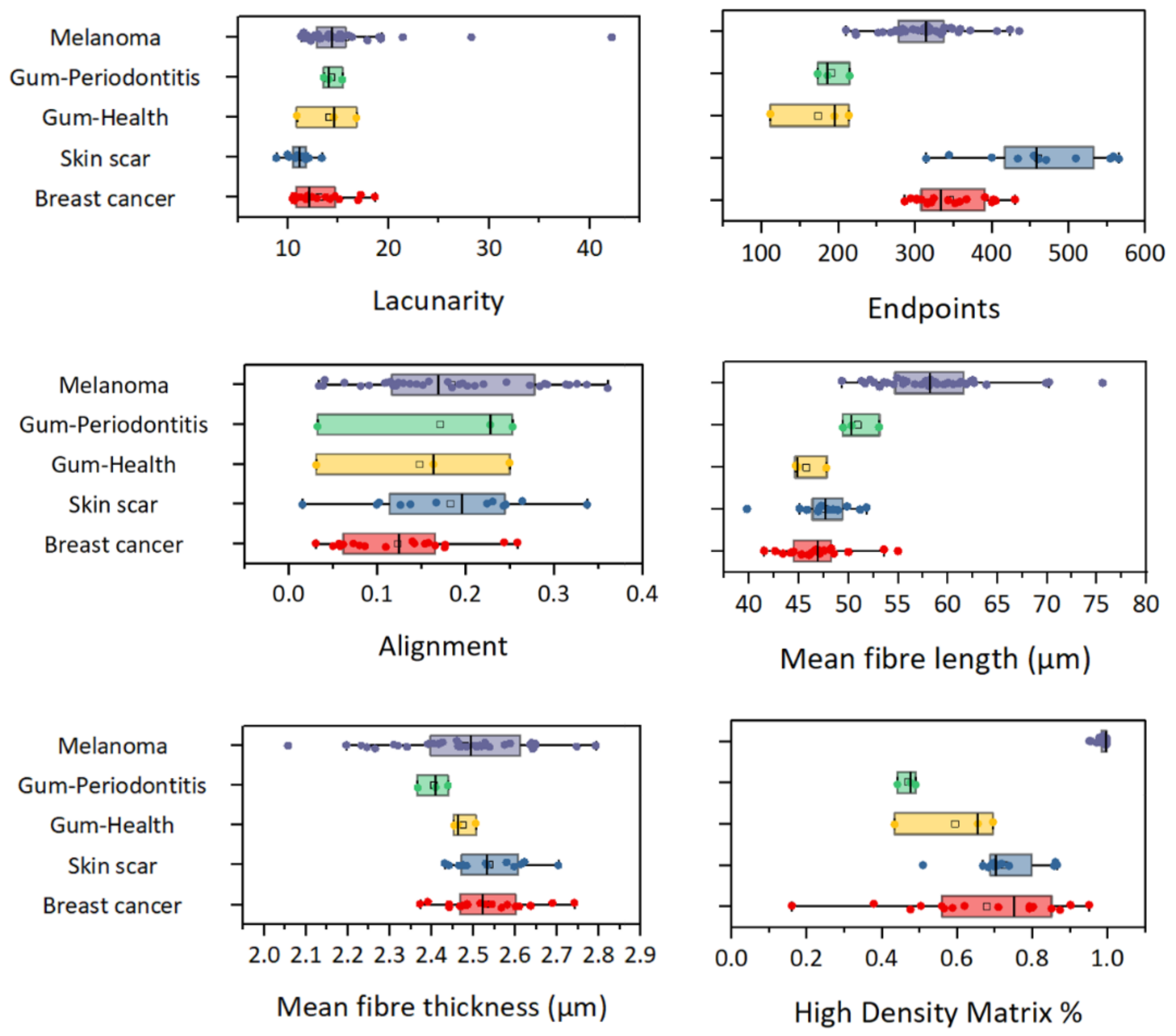


**Supplementary Fig. S5. Quantitative ECM morphometric parameters across pathological and healthy tissues measured by TWOLBMI.** Box-and-whisker plots of six structural descriptors of extracellular matrix (ECM) fiber organization extracted from high-density matrix masks using the TWOLBMI workflow. Tissue types include melanoma (purple), gum–periodontitis (green), gum–healthy (yellow), skin scar (blue), and breast cancer (red). Parameters: Lacunarity (top-left): heterogeneity of pore size distribution within ECM network. Endpoints (top-right): number of fibre termini per unit mask area, reflecting network connectivity. Alignment (middle-left): mean dot-product of local fibre orientation vectors, indicating overall anisotropy. Mean fibre length (middle-right): average continuous segment length of ECM fibres. Mean fibre thickness (bottom-left): average cross-sectional width of fibres. High-density matrix percentage (bottom-right): fraction of the field occupied by high-density matrix pixels. Melanoma and periodontitis ECMs exhibit high lacunarity and endpoint density compared to healthy gum, skin scar, and breast cancer tissues, indicating more fragmented networks. Alignment values vary significantly between diseases, with breast cancer showing the lowest fibre alignment. Box plots show median (center line), quartiles (boxes), and range (whiskers); outliers plotted individually. Color codes correspond to tissue types.

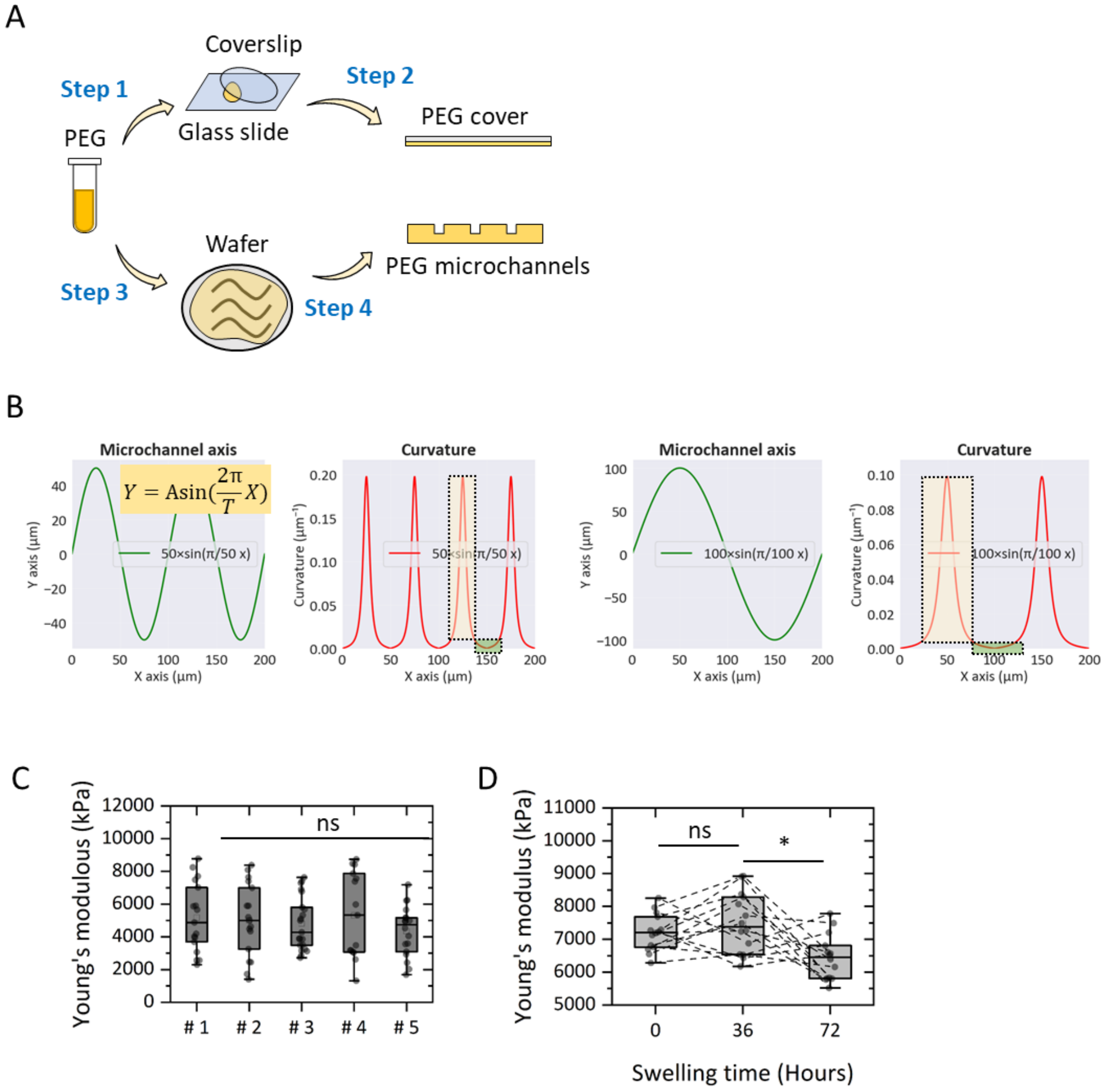


**Supplementary Fig. S6. Fabrication, curvature modelling, and mechanical properties of PEG microchannels.** (**A**) Fabrication workflow: Step 1 — A clean glass coverslip is prepared and surface-coated with PEG. Step 2 — A PEG cover layer is applied. Step 3 — A patterned silicon wafer containing microchannel designs is brought into conformal contact. Step 4 — PEG microchannels are formed via soft lithography and demoulding, yielding sinusoidal profile channels for cell culture experiments. (**B**) Curvature modelling of sinusoidal microchannel geometries with two amplitude-wavelength combinations. Color-coded curvature plots quantify bend magnitude used for high- and low-curvature experimental conditions. (**C**) Young's modulus comparison between five independently fabricated channel batches (#1-#5), showing no statistically significant (ns) differences. #1, n = 15; #2, n = 15; #3, n = 15; #4, n = 15; #5, n = 15. (**D**) Effect of PEG swelling (0 h, 36 h, 72 h in culture medium) on microchannel stiffness; modulus decreased significantly after 72 h, while the 36 h swelling did not differ significantly from baseline. 0-hour, n = 12; 36-hour, n = 15; 72-hour, n = 12. Box plots depict median (center line), quartiles (box edges), and outliers (dots). n.s. indicates non-significant by one-way ANOVA with Tukey post-hoc test.

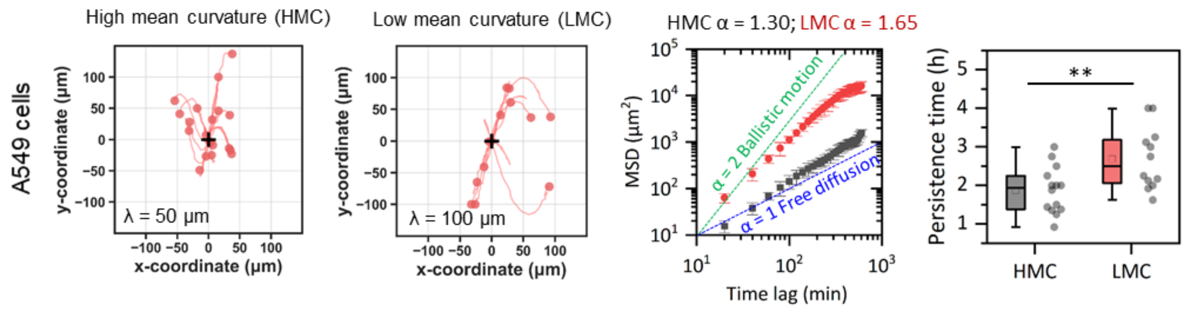


**Supplementary Fig. S7. Extended cell migration analyses in sinusoidal PEG microchannels**. Representative migration tracks within high mean curvature (HMC) and low mean curvature (LMC) sinusoidal channels of A549 cells. Red lines trace individual cell trajectories over the observation period, with the start position marked by a filled circle and the end by an arrowhead; black arrows indicate the overall displacement vector. Mean-square displacement (MSD) analysis (center) shows that trajectories in HMC channels exhibit sub-ballistic motion compared with LMC, with ballistic and free diffusion shown for reference. Persistence time (right) was significantly longer in LMC than HMC, indicating curvature-dependent modulation of persistent migration. HMC, n = 14 cells; LMC, n = 12 cells. Box plots show median and quartiles; statistical significance in panel A assessed by unpaired t-test.

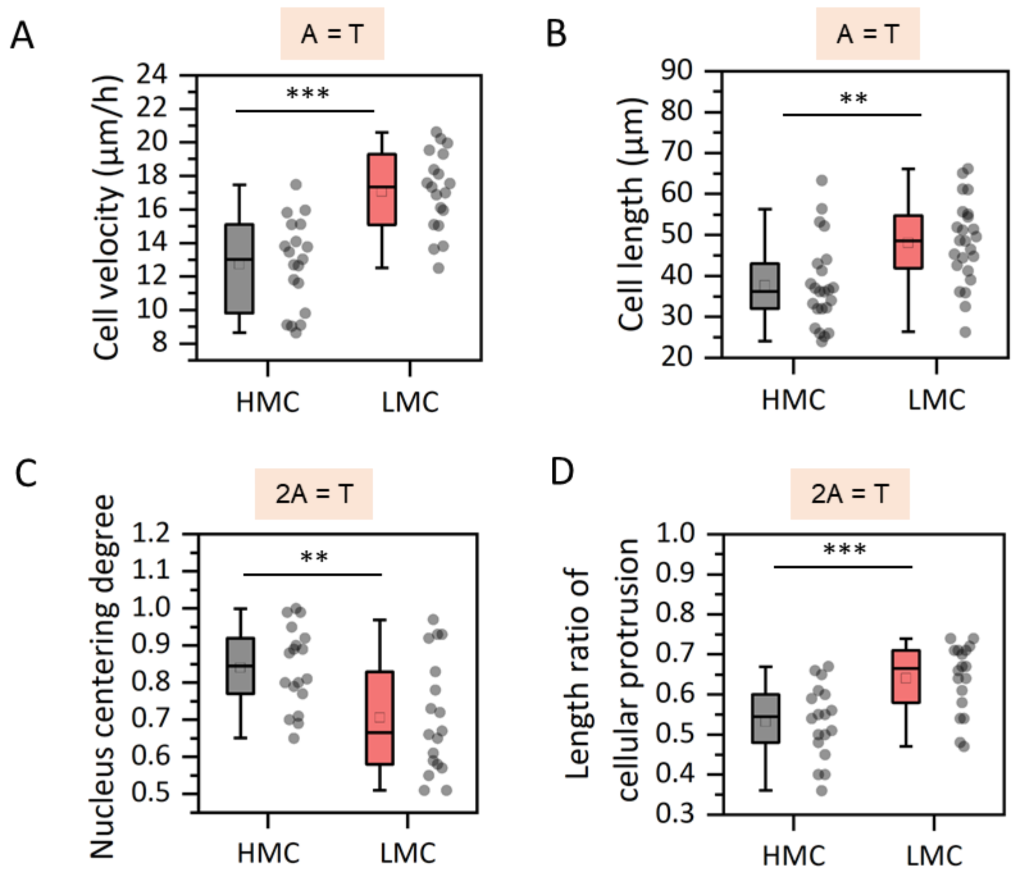


**Supplementary Fig. S8. Influence of channel curvature and geometry parameters on nuclear positioning, protrusion formation, migration speed, and cell length.**

(**A**) Migration speed of A549 cells in channels with 2A = T geometry (amplitude A = half the period T), assessing whether channel wavelength–amplitude relationship influences persistent motility. LMC-like curvature in these channels resulted in significantly faster migration speeds than HMC-like curvature. HMC, n = 18; LMC, n = 18.

(**B**) Cell length in the same 2A = T channel configuration as C. Cells in LMC-like curvature channels were significantly longer than those in HMC-like curvature channels, indicating that reduced curvature facilitates elongation during migration. HMC, n = 18; LMC, n = 18.

(**C**) Nucleus centring degree in A549 cells migrating within high mean curvature (HMC) and low mean curvature (LMC) channels, fabricated using the original geometry configuration (2A ≠ T). HMC conditions yielded significantly higher nuclear centring compared with LMC. HMC, n = 21; LMC, n = 21.

(**D**) Cellular protrusion length ratio (protrusion length relative to total cell length) under the same HMC and LMC conditions as in (A). LMC channels promoted longer relative protrusions than HMC channels. HMC, n = 23; LMC, n = 23. Box plots depict median (center line), quartiles (box edges), range (whiskers). Statistical significance was assessed using unpaired two-tailed Student’s t-tests.

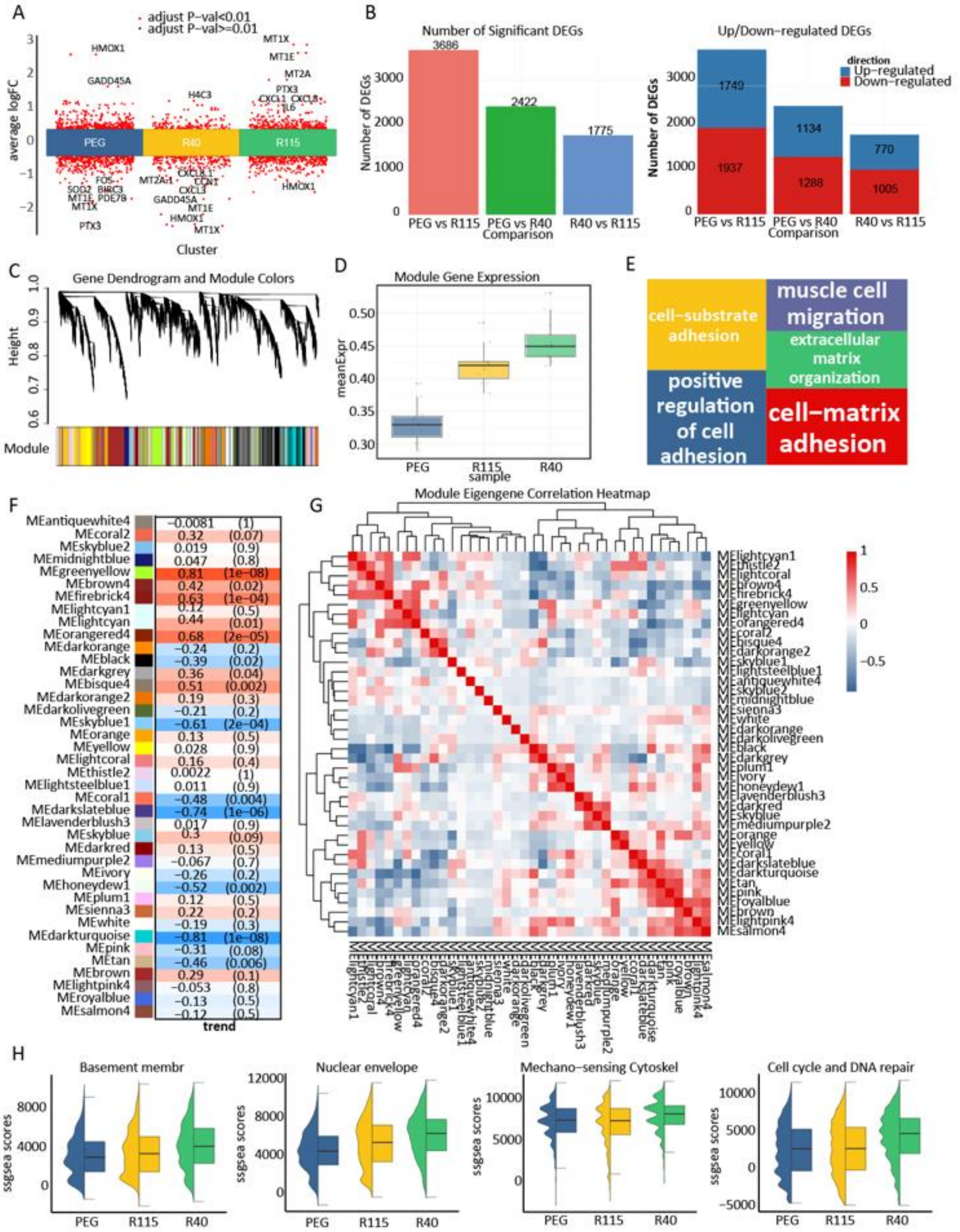


**Supplementary Fig. S9. Transcriptomic modules associated with curvature.**

(**A**) Volcano plot of differentially expressed genes (DEGs) between groups, with curvature-responsive genes highlighted.

(**B**) Number and direction of significant DEGs for each comparison, showing a predominance of upregulated genes under high curvature.

(**C**) Gene dendrogram and module color assignments from WGCNA analysis.

(**D**) Boxplot of light-green module eigengene expression, demonstrating positive correlation with curvature.

(**E**) Functional enrichment of curvature-associated modules, including ECM remodeling and cell–matrix adhesion.

(**F**) Module–curvature correlation heatmap with statistical significance indicated.

(**G**) Module eigengene correlation network showing central positioning of curvature-enriched modules within the co-expression network.

(**H**) Violin plots show upregulation of ssgsea scores for cytoskeleton organization, fibrous crosslinked ECM, basement membrane, DNA repair and nuclear envelope pathways in high-curvature conditions.

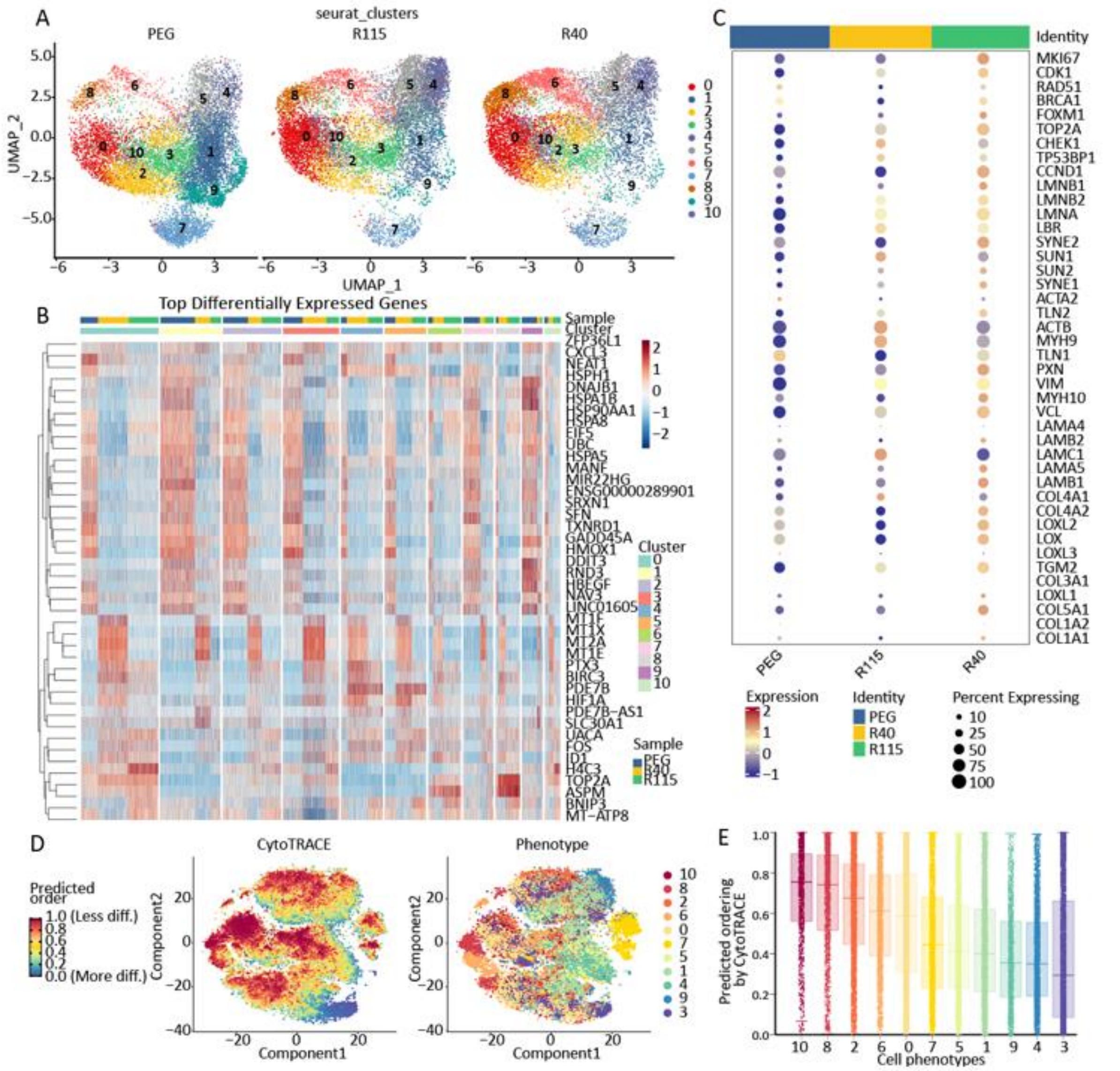


**Supplementary Fig. S10. Single-cell analysis of curvature effects on epithelial subpopulations.**

(**A**) UMAP plots showing curvature-dependent clustering shifts in epithelial subpopulations.

(**B**) Heatmap of top differentially expressed genes across subclusters and curvature conditions.

(**C**) Dot plot displaying expression and fraction of cells expressing key mechanosensitive and ECM-related genes across groups.

(**D**) CytoTRACE analysis indicating lower predicted differentiation potential in curvature-expanded subclusters.

(**E**) Boxplots of predicted differentiation order for each subcluster, with R40-enriched populations showing reduced differentiation scores suggestive of enhanced invasiveness.

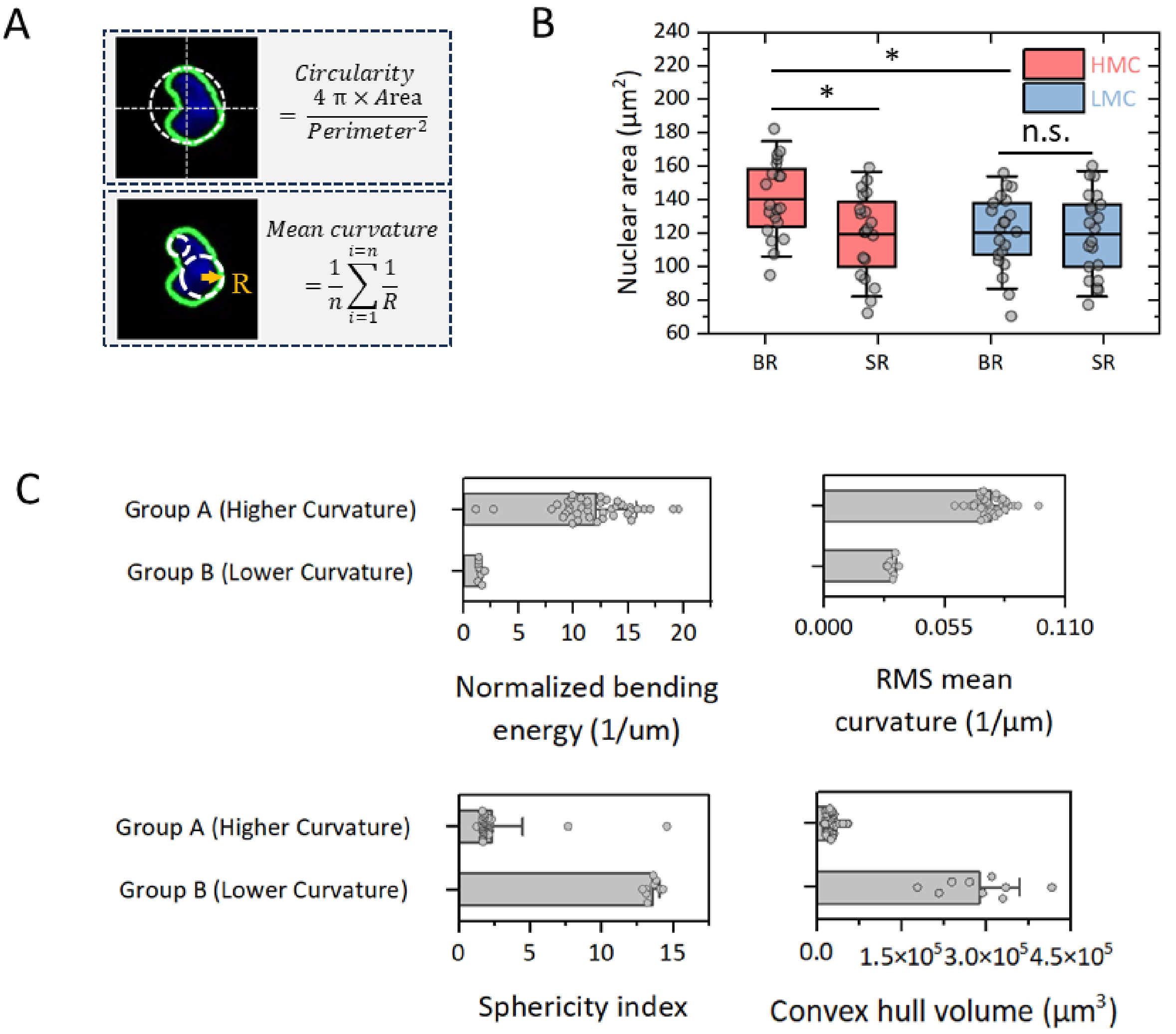


**Supplementary Fig. S11. Nuclear morphometric parameters in cells migrating through sinusoidal PEG microchannels of differing curvature**.

(**A**) Analytical framework for nuclear shape quantification.

(**B**) Nuclear area in A549 cells migrating in high mean curvature (HMC) versus low mean curvature (LMC) microchannels, separated into BR and SR subgroups. HMC channels induced significantly larger nuclear areas in both BR and SR cells compared with LMC channels, with no significant difference between BR and SR within LMC conditions. HMC-BR, n = 20; HMC-SR, n = 20; LMC-BR, n = 21; LMC-SR, n = 21.

(**C**) Extended nuclear morphology metrics: Normalized bending energy (top-left) was markedly higher in Group A nuclei compared with Group B, consistent with increased shape deformation costs. Root-mean-square (RMS) mean curvature (top-right) revealed elevated curvature variance in Group A, indicating more pronounced nuclear bending. Coefficient of variation (CV) of mean curvature (bottom-left) demonstrated greater heterogeneity of local curvature in Group A. Convex hull volume (bottom-right) indicated significant differences in

3D nuclear envelope expansion between groups. Box plots depict median (centre line), quartiles (box edges), range (whiskers), and outliers (dots). Statistical significance was assessed by unpaired two-tailed Student's t-test unless otherwise indicated; n.s., non-significant results.

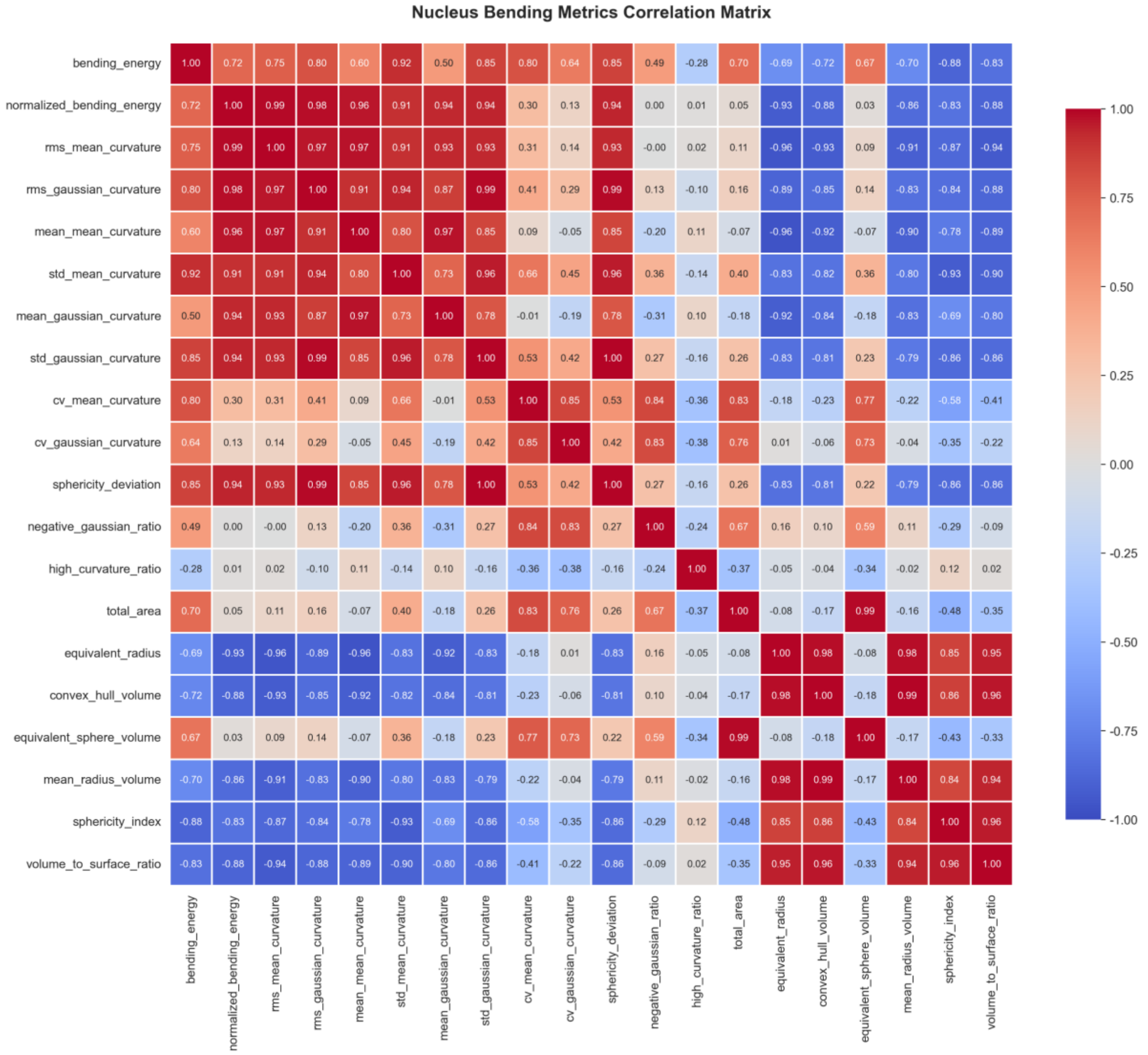


**Supplementary Fig. S12. Correlation matrix of nuclear bending and morphometric parameters**. Pearson correlation coefficients (colour-coded) between 20 nuclear shape descriptors extracted from 3D reconstructions of migrating cell nuclei in sinusoidal PEG microchannels. Warmer colours (red) indicate positive correlation; cooler colours (blue) indicate negative correlation; intensity reflects correlation strength (*r* values from,1 to +1). Metrics include bending energy, normalized bending energy, RMS and mean curvature (both Gaussian and mean curvature estimators), standard deviation and coefficient of variation (CV) of curvature, sphericity deviation, negative Gaussian ratio, high curvature ratio, total nuclear area, equivalent radius, convex hull volume, equivalent sphere volume, mean radius in volume, sphericity index, and volume-to-surface ratio. Strong positive correlations (*e.g.,* bending energy ↔ normalized bending energy, ($r > 0.99$)) reflect mathematically or morphologically linked quantities. Negative associations (*e.g.,* convex hull volume ↔ normalized bending energy) indicate trade-offs between volumetric expansion and bending severity. Parameters such as high curvature ratio show weaker correlations with overall size metrics, suggesting partial independence from nuclear volume changes.

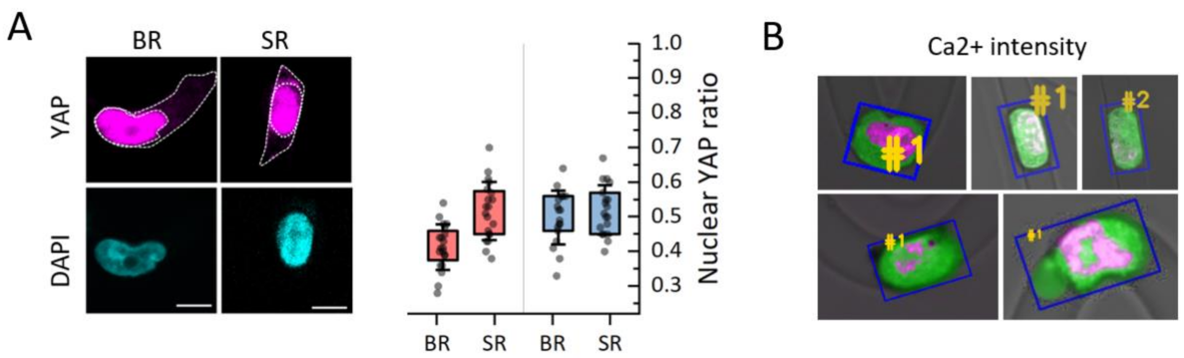


**Supplementary Fig. S13. Nuclear YAP localisation and intracellular $Ca^{2+}$ intensity in bent-run (BR) and straight-run (SR) migrating cells**.

(**A**) Nuclear YAP ratio analysis in A549 cells migrating in sinusoidal PEG microchannels under high mean curvature (HMC; red) or low mean curvature (LMC; blue) conditions, classified by migration conformations: BR, bent trajectories with frequent direction changes, and SR, straight trajectories with minimal heading deviation. Immunofluorescence shows YAP (magenta) and nuclei counterstained by DAPI (cyan). Quantification of nuclear YAP ratio (nuclear YAP intensity / total cellular YAP intensity) revealed no statistically significant differences between BR and SR within each curvature condition (unpaired t-test). Scale bars, 10 µm. HMC-BR, n = 17; HMC-SR, n = 17; LMC-BR, n = 21; LMC-SR, n = 17.

(**B**) Intracellular $Ca^{2+}$ intensity mapping in migrating cells. Live-cell fluorescence imaging was performed using $Ca^{2+}$-sensitive dye (magenta) overlaid on transmitted-light images; blue masks indicate automated cell outline detection, and numbered regions (#1, #2) correspond to nuclear regions and selected cytoplasmic domains used for quantitative analysis. Representative images illustrate heterogeneity of $Ca^{2+}$ distribution between nuclei and surrounding cytoplasm during migration.

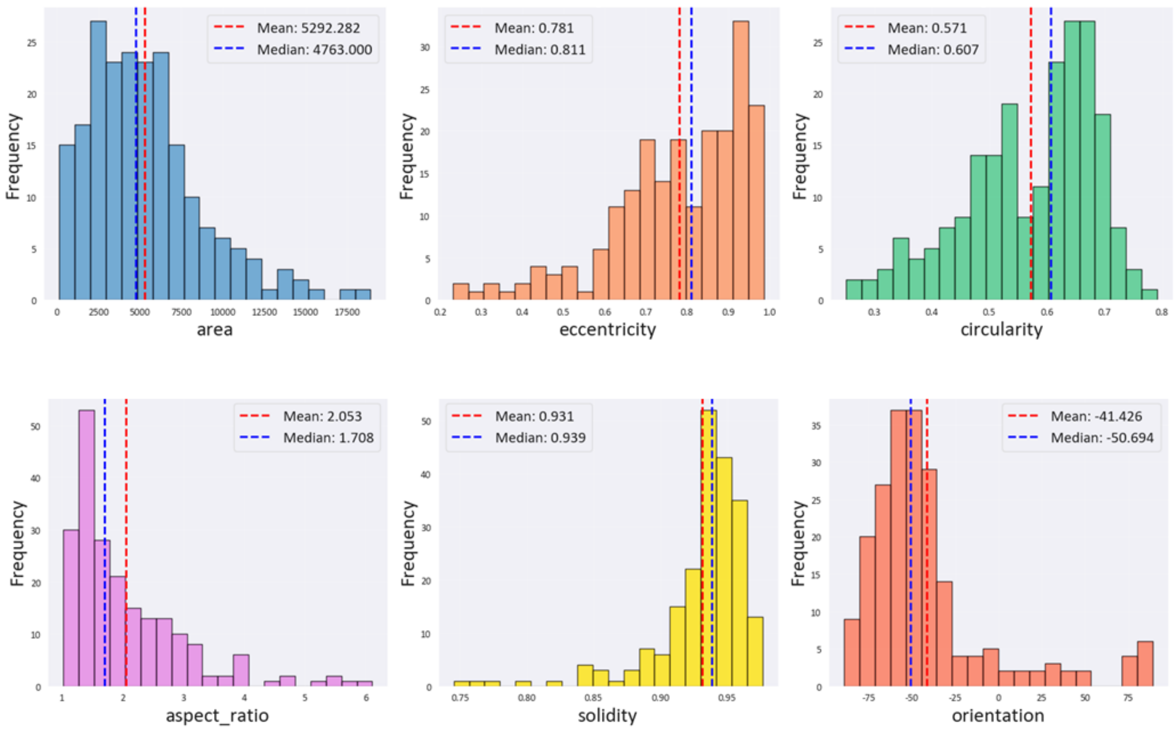


**Supplementary Fig. S14. Distribution of morphometric parameters for nuclei analysed**. Histograms summarising the distribution of six nuclear shape parameters across all analysed cells in sinusoidal PEG microchannels, corresponding to quantitative datasets shown in the main text. Area (pixels) frequency distribution, indicating the prevalence of smaller-to-moderately sized nuclei with a right-tailed spread; vertical dashed lines denote the mean (blue) and median (red) values. Eccentricity distribution (ratio describing long axis extension relative to a perfect circle), showing a bias towards moderately elongated shapes. Circularity distribution, reflecting variability from nearly circular to more irregular nuclear outlines. Aspect ratio distribution (length/width), with peaks at ~1.7-2.0, indicating dominance of mildly elongated nuclear forms and a minority of highly stretched shapes. Solidity distribution (Area / Convex hull area), approaching unity in most cases, consistent with compact nuclear contours and limited concavity. Orientation distribution (principal axis angle relative to channel axis), demonstrating preferential alignment with negative orientation angles in the given experimental setup. Dashed lines indicate the mean (blue) and median (red) of each parameter; summary statistics are provided above each panel. Analysis encompasses all nuclei segmented and measured as described in Methods: Nuclear shape quantification.

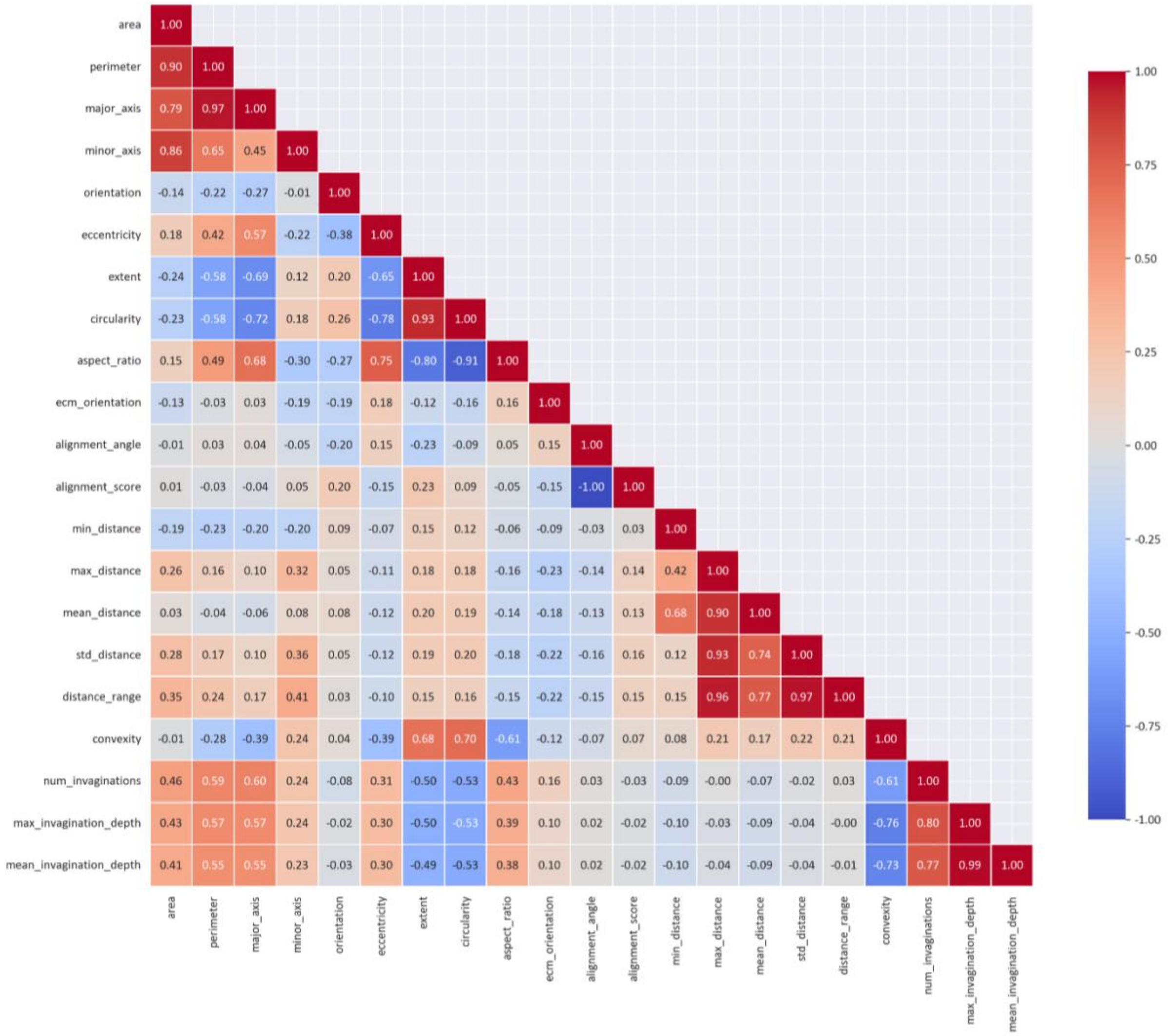


**Supplementary Fig. S15. Correlation matrix of morphometric and invagination metrics for nuclei analysed.** Pearson correlation coefficients (colour-coded) between 22 nuclear morphometric and contour-invagination descriptors extracted from all segmented nuclei in sinusoidal PEG microchannels. Warmer colours (pink/red) denote positive correlations; cooler colours (blue) denote negative correlations; intensity represents correlation strength. Parameters include size metrics (area, perimeter, major/minor axis lengths), shape descriptors (eccentricity, aspect ratio, circularity, extent, orientation, ecn_orientation), alignment measures (alignment_angle, alignment_score), distance-based contour measures (min_distance, max_distance, mean_distance, std_distance, distance_range), compactness (convexity), and nuclear envelope invagination properties (num_invaginations, max_invagination_depth, mean_invagination_depth). Strong positive correlations observed between area and perimeter, major/minor axis lengths, and invagination depth metrics (max_invagination_depth ↔ mean_invagination_depth reflect direct geometrical coupling. Negative correlations (e.g., aspect ratio ↔ circularity; convexity ↔ num_invaginations indicate trade-offs between nuclear elongation, outline smoothness, and discontinuities in contour.

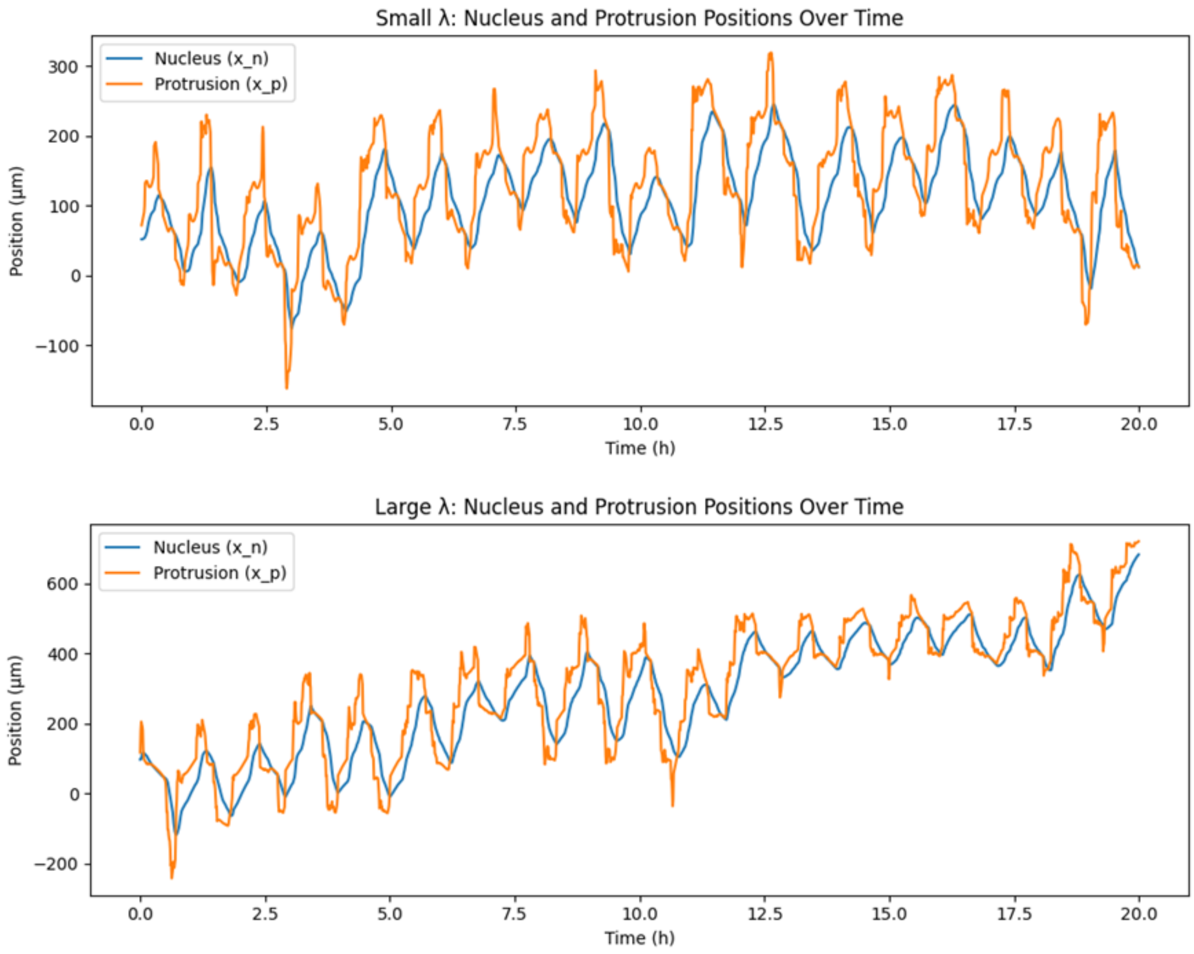


**Supplementary Fig. S16. Theoretical model outputs illustrating the coupling between nuclear and protrusion dynamics under curvature-dependent signalling**. Small wavelength parameter (λ) simulations: temporal trajectories of the nucleus (blue) and leading-edge protrusion (orange) show frequent position fluctuations and phase shifts during migration, consistent with oscillatory behaviour induced by high curvature and short spatial periodicity in the channel geometry. Large wavelength parameter (λ) simulations: nucleus and protrusion positions exhibit sustained forward progression with minimal relative lag, reflecting stable migration states facilitated by low curvature gradients and long spatial periodicity.

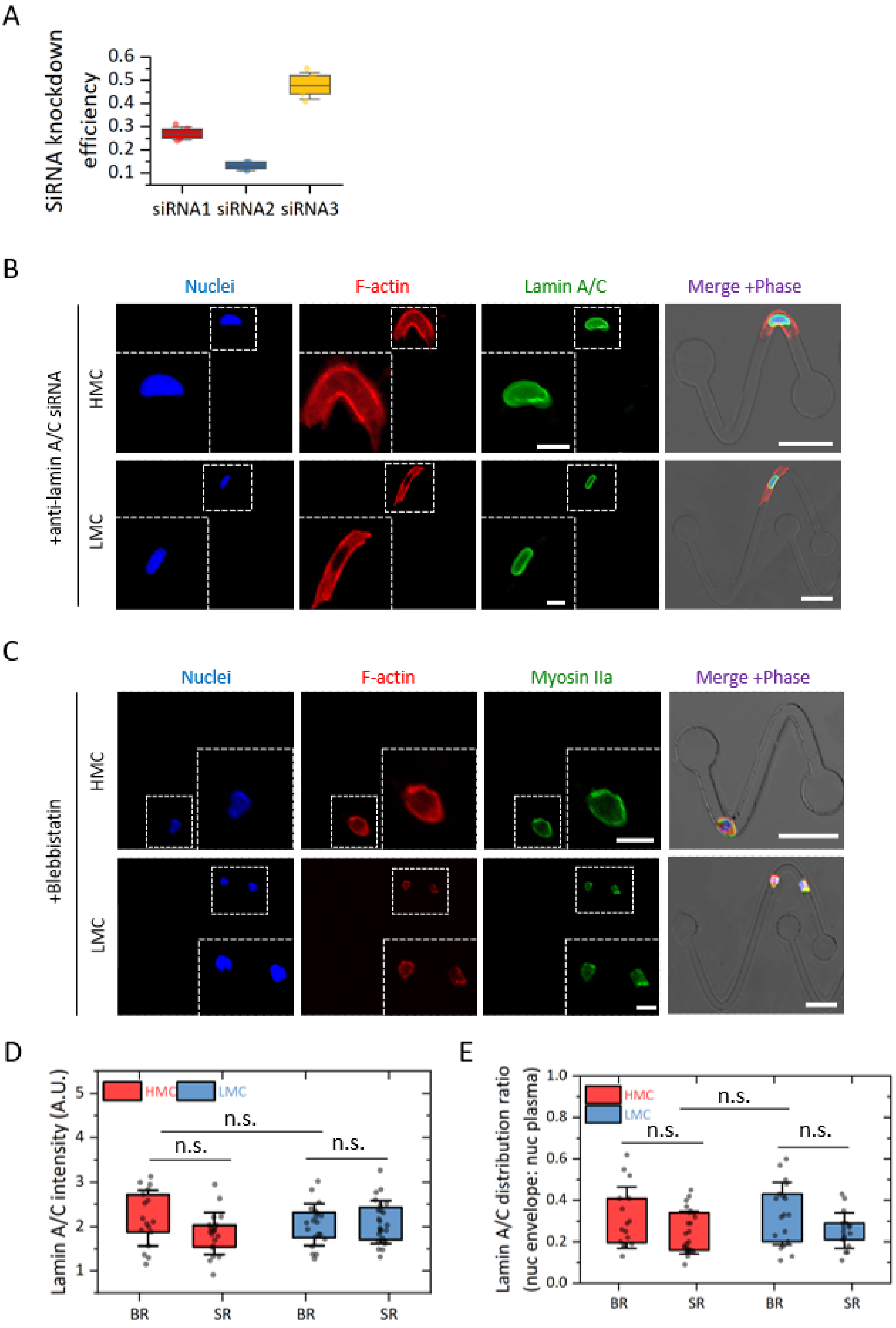


**Supplementary Fig. S17. Effects of Lamin A/C knockdown and myosin inhibition on cortical cytoskeleton organisation in curved microchannels.**

(**A**) siRNA knockdown efficiency for three independent sequences (siRNA1–3) targeting Lamin A/C, expressed from quantitative immunofluorescence measurements. siRNA3 produced the highest knockdown efficiency among the tested constructs.

(**B**) Lamin A/C knockdown disrupts nuclear envelope morphology and peri-nuclear F-actin organisation. Representative fluorescence micrographs show nuclei (DAPI; blue), F-actin (phalloidin; red), Lamin A/C (green), and merged images overlaid on phase contrast in A549 cells migrating through high mean curvature (HMC) or low mean curvature (LMC) sinusoidal microchannels under Lamin A/C suppression. Insets display magnified views of nuclear and cortical regions. Scale bars, 10 μm.

(**C**) Blebbistatin treatment inhibits myosin IIa activity while preserving gross nuclear morphology. Micrographs show nuclei (blue), F-actin (red), myosin IIa (green), and merged phase overlays for cells in HMC and LMC channels following blebbistatin exposure. Insets highlight changes in cytoskeletal architecture. Scale bars, 10 μm.

(**D**) Total Lamin A/C fluorescence intensity in bent-run (BR) and straight-run (SR) cells under HMC or LMC conditions. No statistically significant differences were detected (n.s.). HMC-BR, n = 20; HMC-SR, n = 19; LMC-BR, n = 20; LMC-SR, n = 19.

(**E**) Lamin A/C distribution ratio (mean nuclear envelope intensity / mean nucleoplasmic intensity) across BR and SR subgroups. Ratios were consistent between curvature conditions and migration conformations (n.s.). HMC-BR, n = 20; HMC-SR, n = 19; LMC-BR, n = 20; LMC-SR, n = 19. Statistical comparisons in (D) and (E) employed unpaired two-tailed Student's t-test; box plots show median (centre), quartiles (box) and individual cell measurements (dots).

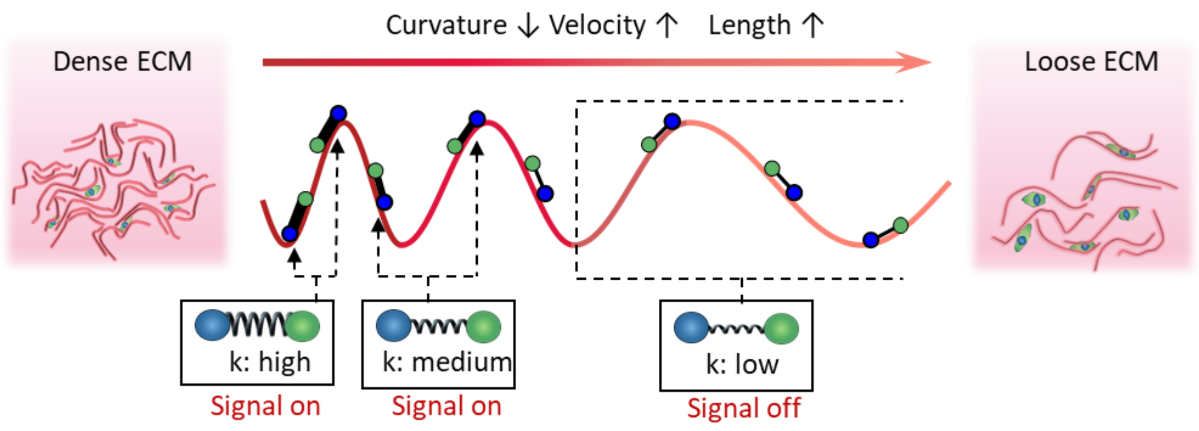


**Supplementary Fig. S18. Conceptual model linking extracellular matrix (ECM) density, curvature modulation, and curvature-dependent signalling.** Schematic illustration showing how sinusoidal trajectories of varying curvature mediate nuclear–protrusion coupling and associated signal activation in migrating cells across heterogeneous ECM densities. At high ECM density (left), cells encounter frequent high-curvature arcs (short wavelength), represented by red sinusoidal traces, which induce repeated mechanical stretching of the nuclear–protrusion linkage. In these regimes, curvature-dependent mechanosensory pathways (*e.g.*, Rac1 activation) remain switched on due to sustained local deformation and force generation. As curvature decreases (centre), average migration velocity increases and path length extends, with only occasional moderate-curvature regions still triggering signal activation. In low ECM density environments (right), long-wavelength trajectories dominate, curvature and mechanical coupling fall below activation thresholds, and mechanosensory signalling switches off. This conceptual framework integrates empirical findings with the proposed curvature-velocity-signalling model and highlights how ECM structural organisation could spatially gate signal transduction via geometry-dependent nuclear-cortex mechanics.

**Table S1. Parameter Table**

| Symbol | Code Parameter | Value | Description |
|---|---|---|---|
| **Geometry** | | | |
| $A$ | `A_small` / `A_large` | 100 / 300 | Amplitude of sinusoidal substrate ($\mu m$). |
| $T$ | `T_small` / `T_large` | 100 / 300 | Period/Wavelength of substrate ($\mu m$). |
| $\kappa_{max}$ | `max_curvature` | 0.002 | Normalization constant for curvature ($1/\mu m$). |
| **Mechanics** | | | |
| $L_0$ | `L0` | 20.0 | Equilibrium cell length ($\mu m$). |
| $k_0$ | `k0` | 0.1 | Basal spring stiffness ($nN/\mu m$). |
| $\alpha_{spring}$ | `a_spring_curv` | 1000.0 | Curvature stiffening coefficient. |
| $F_{max}$ | `F_protru` | 100.0 | Maximum active protrusion force ($nN$). |
| $\lambda$ | `lamda_small/large` | 0.008 | Basal friction coefficient ($nN \cdot h/\mu m$). |
| **Dynamics** | | | |
| $dt$ | `dt` | 0.01 | Time step size (hours or normalized time). |
| $\eta$ | `noise_scale_factor` | 1/150 | Noise scaling factor relative to period $T$. |
| **Decision** | | | |
| $\Delta_{thresh}$ | `turn_threshold` | 0.2 | Rac1-RhoA difference threshold for stability. |
| $P_{base}$ | `turn_base_prob` | 0.1 | Base probability for random turning. |
| $N_{freeze}$ | `turn_freeze_steps` | 50 | Refractory period steps after a turn. |
